# Changes in behavioral priority influence the accessibility but not the quality of working memory content

**DOI:** 10.1101/2022.08.12.503778

**Authors:** Edward F. Ester, Paige Pytel

## Abstract

Evolving behavioral goals require the existence of selection mechanisms that prioritize task-relevant working memory (WM) content for action. Selecting an item stored in WM is known to blunt and/or reverse information loss in stimulus-specific representations of that item reconstructed from human brain activity, but extant studies have focused on all-or-none circumstances that allow or disallow an agent to select one of several items stored in WM. Conversely, behavioral studies suggest that humans can flexibly assign different levels of priority to different items stored in WM, but how doing so influences neural representations of WM content is unclear. One possibility is that assigning different levels of priority to items in WM influences the quality of those representations, resulting in more robust neural representations of high- vs. low-priority WM content. A second – and non-exclusive – possibility is that asymmetries in behavioral priority influence how rapidly neural representations of high- vs. low-priority WM content can be selected and reported. We tested these possibilities in two experiments by decoding high- and low-priority WM content from EEG recordings obtained while human volunteers performed a retrospectively cued WM task. Probabilistic changes in the behavioral relevance of a remembered item had no effect on our ability to decode it from EEG signals; instead, these changes influenced the latency at which above-chance decoding performance was reached. Thus, our results indicate that probabilistic changes in the behavioral relevance of WM content influence the ease with which memories can be accessed and retrieved independently of their strength.

## Introduction

Flexible behaviors require sensory inputs to be compared with internal representations of goal states. Many of these comparisons take place in working memory (WM), a capacity- and duration-limited system that forms a temporal bridge between fleeting sensory phenomena and possible actions (van Ede & Nobre, 2022). Changing environmental circumstances and evolving behavioral goals necessitate the existence of internal selection mechanisms that prioritize task-relevant WM contents for action, especially when an agent must select from among multiple prospective actions or execute a series of actions in sequence (e.g., Cisek & Kalaska, 2010; Cisek, 2019). For example, making your favorite cup of coffee involves performing a series of actions in a specific order, and only when certain external conditions – e.g., the water in the kettle has begun to boil – are met.

The neural consequences of assigning priority to specific WM content can be studied by measuring brain activity linked with WM storage while participants perform a retrospectively cued memory task (Griffin & Nobre, 2003; Lewis-Peacock et al., 2012; Ester et al., 2018; Panichello & Buschman, 2021). In a typical retrocue experiment, an agent remembers an array of items over a brief delay and uses this information to perform a memory-guided behavior. During storage, an informative cue instructs the observer which remembered item(s) are most likely to be required for action at the end of the trial. The typical finding is that an informative retrocue improves memory performance relative to an uninformative cue or no-cue condition (see Souza & Oberauer, 2016, and Myers et al., 2017, for recent comprehensive reviews). Moreover, improvements in memory performance are typically accompanied by improvements in the quality of stimulus-specific WM representations reconstructed from human brain activity, with informative retrocues arresting or even reversing information loss that accumulates during WM storage in the absence of a cue (Sprague et al., 2014; Sprague et al., 2016; Ester et al., 2018; Nouri & Ester, 2020).

With notable exceptions (e.g., Berryhill et al., 2012; Shimi et al., 2014; Günseli et al., 2015; Günseli et al., 2019) most retrocue studies have used perfectly reliable cues. That is, when an informative retrocue appears, it indicates which of a set of remembered items will be later probed with perfect accuracy. Living organisms, however, exist in dynamic natural environments where the future can take on several possibilities. Thus, the likelihood that that any one piece of information stored in WM will become behaviorally relevant is best understood as a matter of probability rather than a certainty. Behavioral studies suggest that retrocue benefits on WM performance are probabilistic and scale with cue reliability; thus, human observers can flexibly assign different levels of behavioral priority to different items stored in memory. However, less is known about how graded changes in behavioral relevance influence neural representations of WM content. One possibility is that probabilistic changes in behavioral relevance could modulate memory strength, for example by facilitating the allocation of attentional gain to neural populations encoding high- vs. low-priority WM content (e.g., Bays & Taylor, 2017). Conversely, a second – and non-exclusive – possibility is that probabilistic changes in behavioral relevance could influence modulate how easily memories can be selected for behavioral read-out (e.g., e.g., Souza et al., 2016). A critical test of these alternatives would offer new insights into how internal selective attention is used to flexibly prioritize task-relevant WM content. Here, we provide this test.

In two experiments, we recorded EEG while human volunteers performed a retrospectively cued spatial WM task. Across experimental blocks, we varied retrocue reliability between 100% (i.e., perfectly predictive) or 75% and quantified how this manipulation influenced our ability to decode remembered information from scalp EEG measurements. Cue reliability had no influence on maximum decoding performance in either experiment; instead, it had a large effect on the latency at which maximum decoding performance was reached. Thus, we show that probabilistic changes in behavioral priority affect the accessibility but not the quality of stimulus-specific WM representations.

## Methods

### Participants

A total of 71 human volunteers (both sexes) were enrolled in this study. 36 participants completed Experiment 1 and 35 participants completed Experiment 2. Sample sizes in each experiment are commensurate – if not slightly larger – than prior work using similar experimental and analytic approaches (e.g., Wolff et al., 2017; Ester et al., 2018; Ester & Nouri, 2020). Participants were recruited from the Florida Atlantic University community and completed a single 2.5-hour testing session in exchange for monetary remuneration ($15/h in amazon.com gift cards). All participants self-reported normal or corrected-to-normal visual acuity and color vision. All study procedures were approved by the local institutional review board, and all participants gave both written and oral informed consent prior to enrolling in the study. Two participants in Experiment 1 and two participants in Experiment 2 voluntarily withdrew from the study prior to completing both cue conditions (i.e., 100% valid vs. 75% valid); data from these participants were excluded from final analyses. Thus, the data reported here reflect 34 participants in Experiment 1 and 33 participants in Experiment 2.

### Testing Environment

Participants were seated in a dimly-lit and acoustically shielded recording chamber for the duration of the experiment. Stimuli were generated in MATLAB and rendered on a 17’’ Dell CRT monitor cycling at 85 Hz (1024 x 768 pixel resolution) via PsychToolbox-3 software extensions (Kleiner et al., 2007). Participants were seated approximately 65 cm from the display (head position was unconstrained). To combat fatigue and/or boredom, participants were offered short breaks at the end of each experimental block.

### Experiment 1 – Spatial Postcue Task

A task schematic is shown in Figure 1. Each trial began with an encoding display lasting 500 ms. The encoding display contained two colored circles (blue and red; subtending 1.75 degrees visual angle [DVA] from a viewing distance of 65 cm) rendered at of eight polar locations (22.5° to 337.5° in 45° increments) along the perimeter of an imaginary circle (radius 7.5 DVA) centered on a circular fixation point (subtending 0.25 DVA) rendered in the middle of the display. The locations of the two discs were counterbalanced across each task (i.e., 100% valid vs. 75% valid), though not necessarily within an experimental block. Participants were instructed to maintain fixation and refrain from blinking for the duration of each trial. The encoding display was followed by a 2.0 sec postcue display. During informative cue trials the fixation point changed colors from black to either blue or red (i.e., matching the color of a remembered disc), while during neutral trials the fixation point changed colors from black to purple (e.g., the “average” of blue and red). The postcue display was followed by a probe display containing a blue or red fixation point, mouse cursor, and question mark symbol (the initial position of the mouse cursor was always on top of the fixation point). Participants were required to click on the location of the disc matching the color of the fixation point. Participants were instructed to prioritize accuracy over speed, but a 2.5 sec response deadline was imposed. The trial ended when participants clicked on a location or the response deadline elapsed. Sequential trials were separated by a 1.5-2.5 sec blank period (randomly and independently selected from a uniform distribution after each trial). An equal number of informative and neutral cue trials was presented during each experimental block.

**Figure 1.**
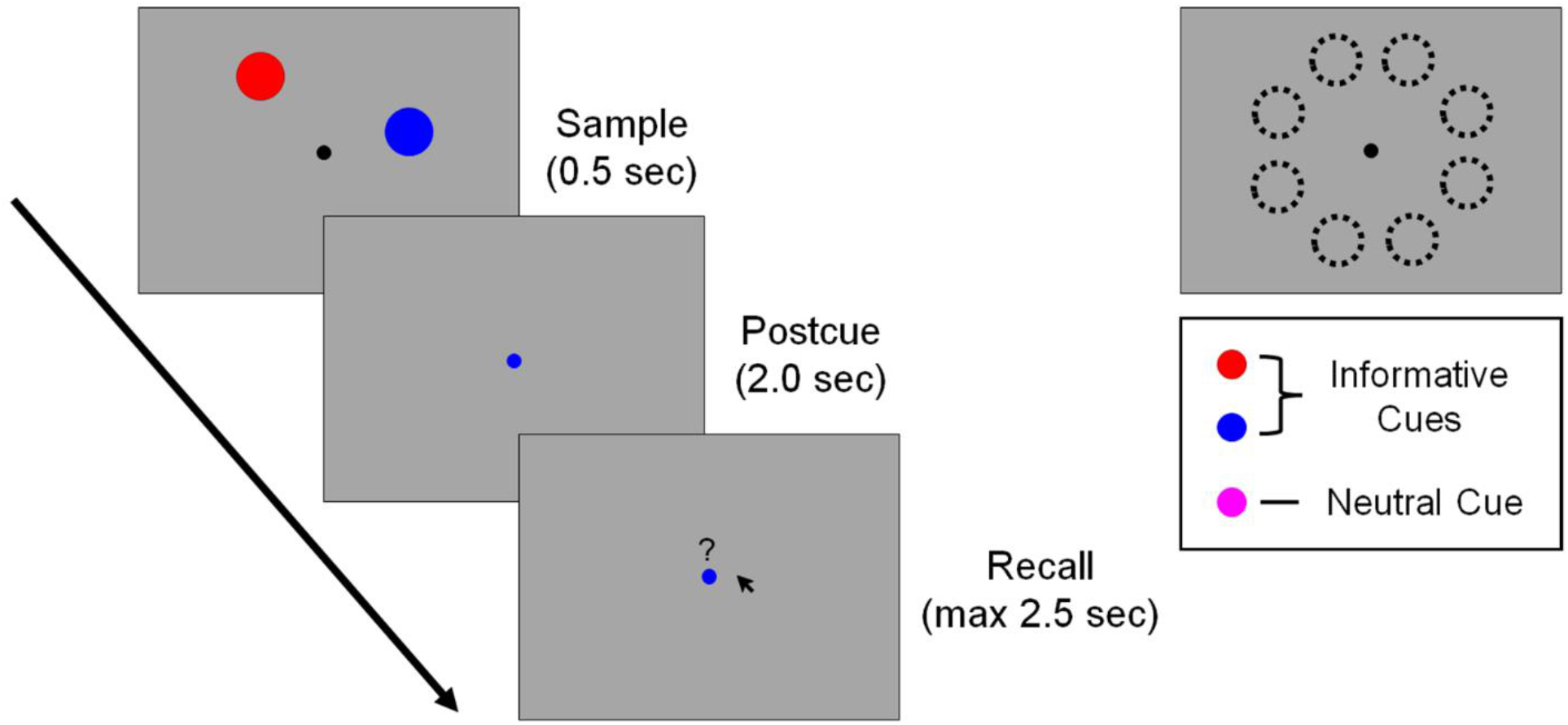
Spatial Postcue Task. Participants encoded the positions of two discs into memory. A postcue instructed participants to remember one disc (informative trials) or both discs (neutral trials) over a blank delay. Participants then recalled the position of the disc matching the color of the fixation point. Discs always appeared in two of eight possible positions (upper right), with disc positions counterbalanced across the experiment. During the first half of experimental blocks postcue reliability was fixed at 100%; during the second half of experimental blocks reliability was lowered to 75%. See text for additional details. Note: the displays shown here are included for exposition and not drawn to scale; see Methods for detailed information regarding stimulus and display geometry.

Each participant completed 12-18 blocks of 56 trials in this task. We had no a priori reason to expect that task order would influence participants’ performance, thus all participants completed the 6-9 blocks of the 100% cue reliability condition followed by 6-9 blocks in the 75% cue reliability condition (the exact number of blocks depended on the time available for testing). During valid cue trials the fixation point remained the same color during the postcue and probe displays; during invalid cue trials the fixation point changed from one color during the postcue display (e.g., blue) to a different color during the probe display (e.g., red). In both cases, participants were instructed to click on the location of the disc matching the color of the fixation point during the probe period.

### Experiment 2 – Spatial Retrocue Task

Experiment 2 was identical to Experiment 1, with the exception that the postcue was replaced by a retrocue presented midway through the delay period (i.e., 1.0 sec after termination of the sample display). We use the term postcue in Experiment 1 and retrocue in Experiment 2 to emphasize that in the former experiment participants can consult an iconic representation of disc positions while in the latter experiment they cannot (Souza & Oberauer, 2016). Like Experiment 1, all participants completed the 6-9 blocks of the 100% cue reliability condition followed by 6-9 blocks in the 75% cue reliability condition (the exact number of blocks depended on the time available for testing).

### Postcues vs. Retrocues

Following earlier work (e.g., Souza & Oberauer, 2016) we use the term postcue to describe circumstances where the timing of an informative cue could allow an observer to consult an iconic representation of to-be-remembered information and the term retrocue to describe circumstances where the timing of an informative cue precludes access to iconic representations. On the assumption that iconic representations tend to persist for ∼300-700 ms (e.g., Sperling, 1960), we refer to informative cues in Experiment 1 – which appear immediately after termination of the sample display – as postcues and informative cues in Experiment 2 – which appear 1.0 sec after termination of the sample display – as retrocues. Some evidence suggests that postcues and retrocues have complementary effects on the quality of WM content. For example, in an earlier study we tracked the quality of position-specific WM representations reconstructed from human EEG activity and found that an informative postcue blunted gradual declines in the quality of WM content while an informative retrocue partially reversed earlier declines in the quality of WM content (Ester et al., 2018). Experiments 1 and 2 in the current manuscript were designed to replicate these findings while also exploring the effects of behavioral priority on the overall quality of position-specific WM representations.

### EEG Acquisition and Preprocessing

Continuous EEG was recorded from 63 uniformly distributed scalp electrodes using a BrainProducts “actiCHamp” system. The horizontal and vertical electrooculogram (EOG) were recorded from bipolar electrode montages placed over the left and right canthi and above and below the right eye, respectively. Online EEG and EOG recordings were referenced to a 64^th^ electrode placed over the right mastoid and digitized at 1 kHz. All data were later re-referenced to the algebraic mean of the left- and right mastoids, with 10-20 site TP9 serving as the left mastoid reference. Data preprocessing was carried out via EEGLAB software extensions (Delorme & Makeig, 2004) and custom software. Data preprocessing steps included the following, in order: (1) resampling (from 1 kHz to 250 Hz), (2) bandpass filtering (1 to 50 Hz; zero-phase forward- and reverse finite impulse response filters as implemented by EEGLAB), (3) epoching from -1.0 to +5.0 relative to the start of each trial, (4) identification, removal, and interpolation of noisy electrodes via EEGLAB software extensions, and (5) identification and removal of oculomotor artifacts via independent components analysis via EEGLAB. After preprocessing, our analyses focused exclusively on the following 10-20 occipitoparietal electrodes: P7, P5, P3, Pz, P2, P4, P6, P8, PO7, PO3, POz, PO2, PO4, PO8, O1, O2, Oz.

### Data Cleanup

Prior to analyzing participants’ behavioral or EEG data, we excluded all trials where the participant responded with a latency of < 0.3 sec (we attributed these trials to accidental mouse clicks following the onset of the probe display rather than a deliberate recall of a specific stimulus position) or failed to respond within the 2.5 sec deadline. This resulted in an average loss (±1 S.E.M.) of 0.429 ± 0.09% trials in Experiment 1 and 0.841 ± 0.22% of trials in Experiment 2.

### Decoding Spatial Positions from Posterior Alpha-Band EEG Signals

Location decoding was based on the multivariate distance between EEG activity patterns associated with memory for specific positions. This approach is an extension of earlier parametric decoding methods (Wolff et al., 2017) designed for use in circular feature spaces. Following earlier work (e.g., Ester et al., 2018), we extracted spatiotemporal patterns of alpha-band activity (8-13 Hz) from 17 occipitoparietal electrode sites (see *EEG Acquisition and Preprocessing* above). The raw timeseries at each electrode was bandpass filtered from 8-13 Hz (zero-phase forward-and-reverse filters as implemented by EEGLAB software), yielding a real-valued signal f(t). The analytic representation of f(t) was obtained via Hilbert transformation:

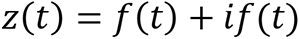

where *i* is the imaginary operator and 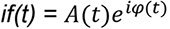. Alpha power was computed by extracting and squaring the instantaneous amplitude A(t) of the analytic signal z(t).

Location decoding performance was computed separately for each disc (i.e., blue vs. red), trial type (i.e., informative vs. neutral) and each task (i.e., 100% vs. 75%) on a timepoint-by-timepoint basis. In the first phase of the analysis, we sorted data from each condition into 5 unique training and test data sets using stratified sampling while ensuring that each training set was balanced across remembered positions (i.e., we ensured that each training data set contained an equal number of observations where the location of the remembered stimulus was at 22.5°, 67.5°, etc.). We circularly shifted the data in each training and test data set to a common center (0°, by convention) and computed trial-averaged patterns of responses associated with memory for each disc position in each training data set. Next, we computed the Mahalanobis distance between trial-wise activation patterns in each test data set with position-specific activation patterns in the corresponding test data set, yielding a location-wise set of distance estimates. If scalp activation patterns contain information about remembered positions then distance estimates should be smallest when comparing patterns associated with memory for identical positions in the training and test data set and largest when comparing opposite positions (i.e., those ±180° apart), yielding an inverted gaussian-shaped function. Trial-wise distance functions were averaged, sign-reversed, and convolved with a cosine function to yield a single decoding estimate for condition and time point with chance decoding yielding a value of 0. Decoding results from each training- and test-data set pair were averaged (thus ensuring the internal reliability of our approach), yielding a single decoding estimate per participant, timepoint, and task condition.

### Quantifying Peak Decoding Accuracy

To determine whether changes in cue reliability influenced the strength of location-specific representations stored in WM, we calculated peak decoding accuracy during 100% reliable and 75% reliable blocks. Peak decoding estimates were then compared with a bootstrap test. We first selected (with replacement) and averaged time courses of decoding activity for the probed location from *N* of *N* participants. Next, we calculated the time of maximum decoding performance following the onset of the probe display (i.e., 2.5 sec after the start of each trial). We defined a 100 ms window around peak decoding performance (i.e., 50 ms before peak decoding performance to 50 ms after it) and used this window to compute temporally averaged peak decoding performance in the same sample of *N* participants. These calculations were performed separately for data from 100% valid and 75% valid blocks and permuted 10,000 times, with a new subsample of participants chosen on every permutation. Finally, we computed the average and 95% confidence interval of peak decoding performance during 100% valid and 75% valid blocks. Statistical significance was assessed by counting the proportion of permutations where peak decoding performance was larger during 100% valid blocks compared to 75% valid blocks.

### Quantifying Peak Decoding Latency

To determine whether changes in cue reliability influenced the timing of WM retrieval, we compared the latencies of above-chance decoding performance during the cue and probe displays. During neutral trials, we computed a cross-correlation between average probe-matching decoding performance from 0.0 to 1.5 sec after the onset of the probe display. Specifically, we calculated the normalized correlation coefficient between the time course of decoding performance during the 100% and 75% blocks while temporally shifting the latter relative to the former by -1.0 to +1.0 sec in 4 ms increments. If time courses of decoding performance during 100% blocks and 75% blocks are identical, then the maximum cross-correlation should be observed at a temporal lag of 0.0 sec. Conversely, if the time-course of above-chance decoding performance during 100% valid blocks precedes the time-course of above-chance decoding performance during 75% valid blocks, then the maximum cross-correlation should occur at a negative temporal lag (i.e., when the time course of decoding performance during 75% blocks is shifted earlier in time). We compared the observed cross correlation function with a null distribution of cross-correlation function estimated by shuffling participant level condition labels (i.e., 100% valid vs. 75% valid) 10,000 times. An analogous approach was used to quantify temporal lags in decoding performance during informative cue trials (whether valid or invalid), with the exception that we used a window spanning -0.5 to 1.5 sec relative to the onset of the probe display. A broader window was deliberately selected as we anticipated cue-matching decoding performance to exceed chance levels during the post- or retrocue period when an informative cue was present.

### Statistical Comparisons – Behavioral Data

Participants’ behavioral data (i.e., absolute average recall error and average response time) were analyzed using standard repeated-measures parametric statistics (e.g., t-test, ANOVA); for these comparisons we report test statistics, p-values, and effect size estimates. Note: given that average absolute recall error can be influence by both memory precision and memory errors (e.g., trials where the participant forgets the location of the probed disc and guesses randomly), we also considered quantifying participants’ memory performance via a two-parameter mixture model which assumes that participants (a) report the location of the probed disc with precision σ or (b) guess randomly with frequency *p* (e.g., Zhang & Luck, 2008). However, preliminary analyses revealed that across all experimental conditions (i.e., valid, neutral, or invalid 100% or 75% trials) random guesses made up a percentage of < 1% of all responses. Thus, we are confident that average absolute recall error estimates reflect the precision of participants’ memory independently of the likelihood of successful recall.

### Statistical Comparisons – EEG Data

The decoding analysis we used assumes chance-level decoding performance of 0. Likewise, direct comparisons of decoding performance or reconstruction strength across conditions (e.g., 100% vs. 75%) assume a null statistic of 0. Thus, we evaluated decoding performance by generating null distributions of decoding performance (or differences in decoding performance across conditions) by randomly inverting the sign of each participant’s data with 50% probability and averaging the data across participants. This procedure was repeated 10,000 times, yielding a 10,000-element null distribution for each time point. Finally, we implemented a cluster-based permutation test (Maris & Oostenveld, 2007; Wolff et al. 2017) with cluster-forming and cluster-size thresholds of p < 0.05 to correct for multiple comparisons across time points. Differences in peak decoding accuracy were quantified with bootstrap tests (see *Quantifying Peak Decoding Accuracy)*, and differences in decoding latency were quantified via randomization tests (see *Quantifying Peak Decoding Latency*)

## Results

### Experiment 1 – Postcues

#### Behavioral Performance

Participants’ memory performance was quantified via average absolute recall error (i.e., the angular difference between the polar location of the probed stimulus and the polar location reported by the participant) and average response latency. During 100% blocks, participants received a neutral or perfectly informative postcue. Conversely, during 75% blocks participants received a neutral, valid, or invalid postcue. Thus, we initially analyzed data from 100% and 75% blocks separately. Participants’ recall errors during 100% blocks and 75% blocks are summarized in Figure 2A. During 100% blocks, participants recall errors were significantly lower during valid relative to neutral trials [t(33) = 4.178, p < 0.0002, d = 0.474]. During 75% blocks, a repeated-measures analysis of variance (ANOVA) with cue type (neutral, valid, or invalid) as the sole factor revealed a main effect, [F(2,66) = 4.456, p = 0.015, η^2^ = 0.119]. To facilitate performance comparisons across different levels of cue reliability (i.e., 100% vs. 75%) we calculated within-condition cue effects by

(a) computing differences in recall error during 100% valid and 100% neutral trials and
(b) computing differences in recall error between 75% valid, 75% neutral, and 75% invalid trials (Figure 2B). Direct comparisons of cue effects during 100% valid and 75% valid blocks indicated that valid cues lowered recall errors by equal amounts during 100% and 75% valid blocks [M = -0.699° and -0.706°, respectively, t(33) = 0.038, p = 0.969]. Conversely, we found no evidence for an invalid cue effect during 75% valid blocks; if anything, recall errors were marginally lower during invalid relative to neutral trials [M = 9.183° vs. 9.225°; t(33) = 0.166, p = 0.869].

**Figure 2.**
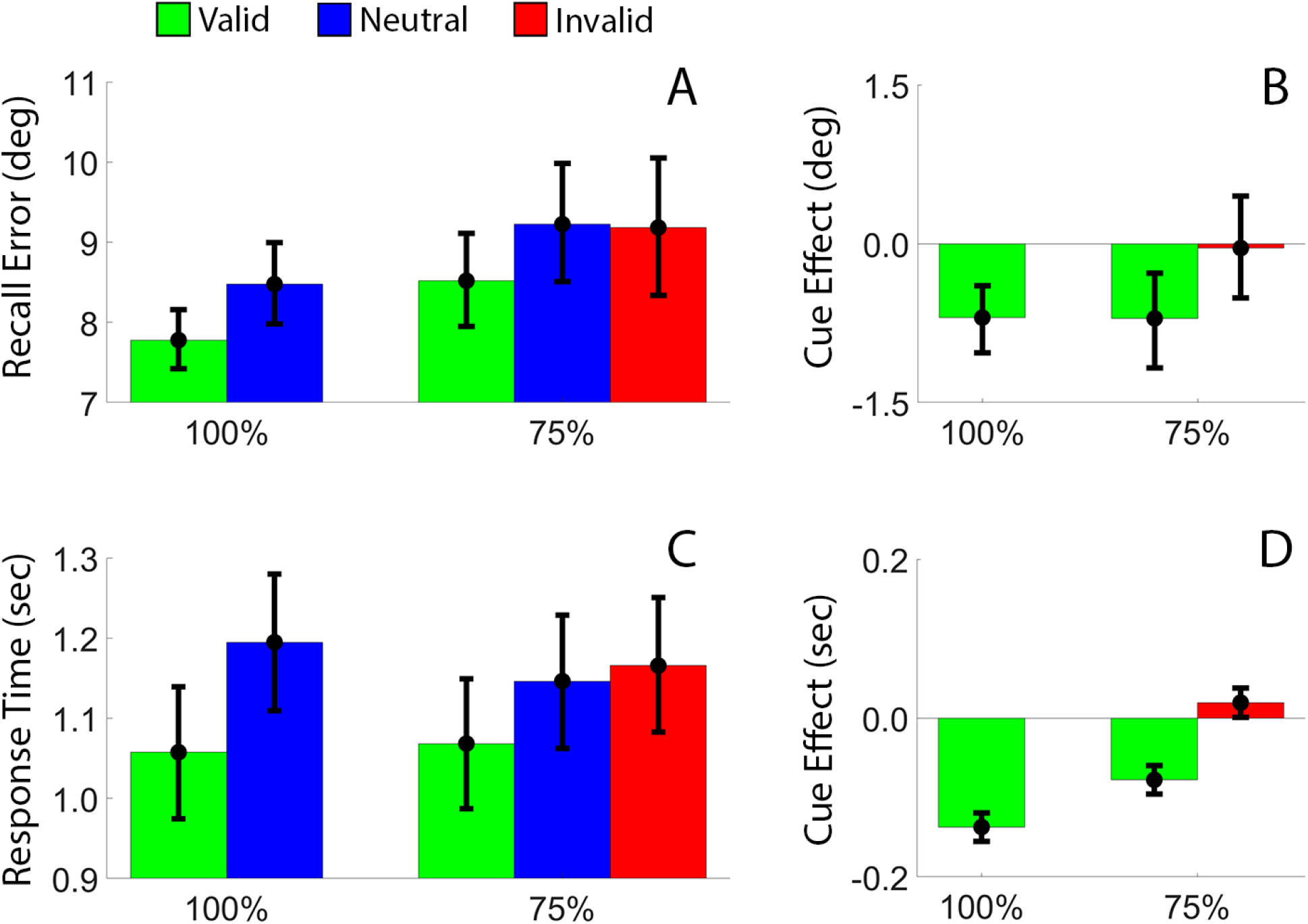
Memory Performance in Experiment 1. (A) Average absolute recall error as a function of cue type (valid, neutral, invalid) and cue reliability (100%, 75%). (B) Cue effects, defined as the difference between valid/invalid and neutral trials. (C) Average response times and (D) cue effects. Error bars depict the 95% confidence interval of the mean.

A complementary analysis of participants’ response times (Figure 2C) revealed a facilitatory effect of valid vs. neutral cues during 100% blocks [t(33) = 14.540, p < 1e-16, d = 0.543], and a repeated measures ANOVA revealed a significant effect of cue type during 75% blocks [F(2,66) = 46.749, p < 1e-14; η^2^ = 0.586]. Analyses of cue effects (Figure 2D) revealed a significantly larger effect of valid cues on response times during 100% vs. 75% blocks [M = 0.136 vs. 0.078 sec, respectively; t(33) = 5.884, p < 1.35e-6, d = 1.088]. Once again, we found no evidence for an invalid cue cost during 75% valid blocks (M = 19 ms; t(33) = 2.006, p = 0.0532, d = 0.078). Thus, valid cues presented during 100% and 75% blocks led to equal improvements in recall performance and parametric improvements in response times compared to neutral cues. Conversely, we found no evidence indicating that invalid cues impaired either recall performance or response times.

#### EEG Decoding Performance

We used a decoding analysis to quantify how changes in behavioral priority influenced location-specific representations stored in WM. During neutral trials, an uninformative postcue instructed participants to remember the positions of both discs presented in the sample display; 2.0 sec later a probe display prompted participants to report the location of one disc via a mouse click. Based on earlier findings (e.g., Ester et al., 2018; Nouri & Ester, 2019; Ester & Nouri, 2022) we expected equivalent decoding performance for each disc during the sample and postcue displays, but significantly higher decoding performance for the probed relative to unprobed disc during the probe display. Moreover, we expected equivalent performance across different levels of cue reliability, i.e., 100% vs. 75% blocks. These predictions were borne out in analyses of location decoding performance during neutral trials (Figure 3). Decoding performance for the disc that was ultimately probed and the disc that was not ultimately probed increased rapidly during the sample display but fell back to chance levels during the ensuring postcue display. Decoding performance for the probe-matching disc – but not the probe-nonmatching disc – increased rapidly following onset of the probe display before returning to chance levels shortly after participants responded. Nearly identical patterns of decoding performance were observed during 100% valid blocks (Figure 3A) and 75% valid blocks (Figure 3B), These findings were expected and are a straightforward replication of earlier results (Ester et al., 2018; Nouri & Ester, 2020).

**Figure 3.**
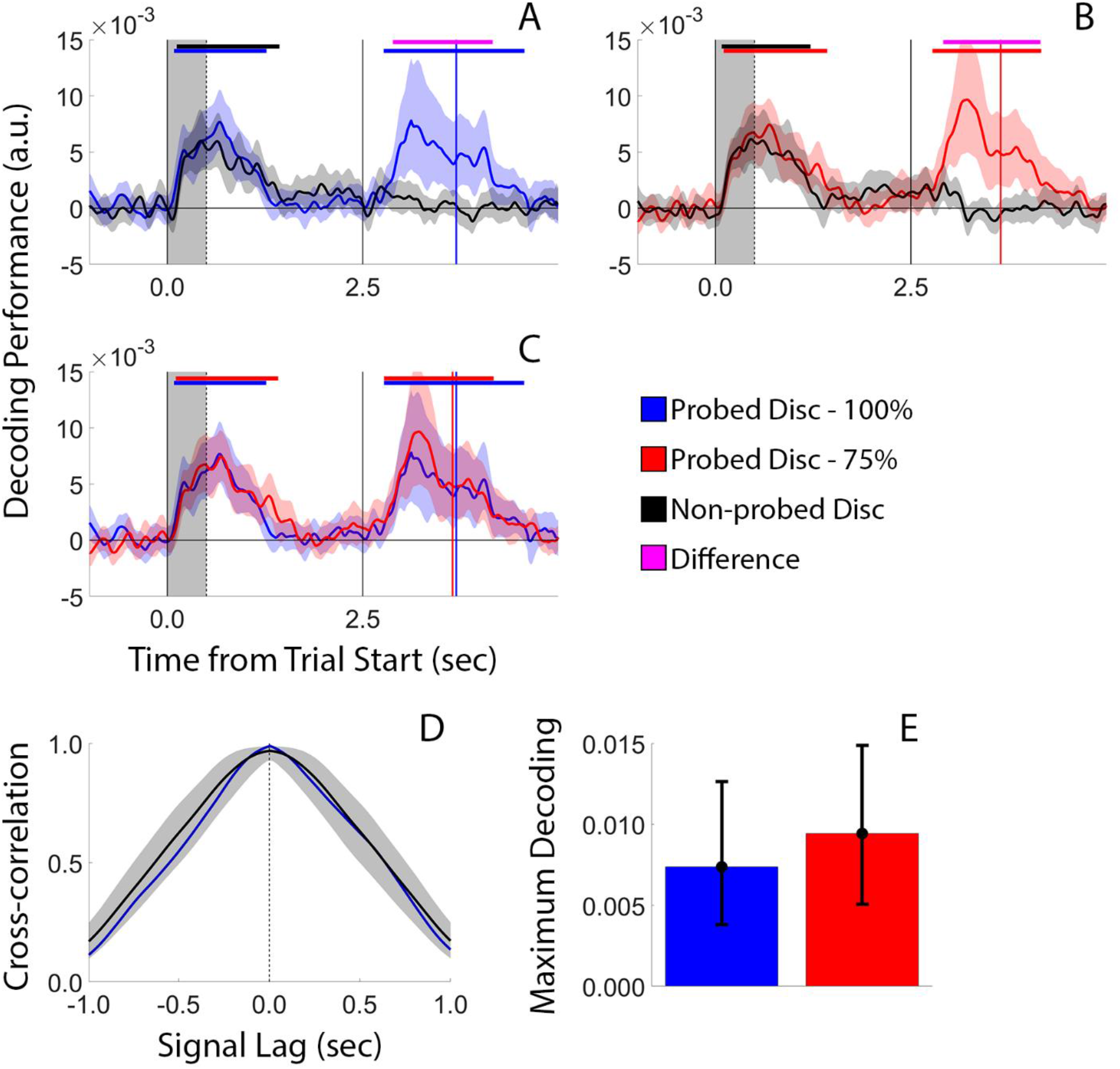
Decoding Performance During Neutral Trials. (A) Average decoding performance for the probe-matching and probe-nonmatching discs during neutral trials of 100% reliable blocks. (B) Identical to panel A, but for neutral trials of 75% blocks. (C) Overlay of probe-matching decoding performance from 100% reliable and 75% blocks (i.e., blue traces in panels A and B). The grey shaded area in each plot marks the sample display; vertical lines at time 0.0, 0.5, and 2.5 mark the onset of the sample, postcue, and response displays, respectively. Blue and red vertical lines mark the average response time across participants. Horizontal bars at the top of each plot mark epochs where decoding performance was significantly > 0 or epochs where decoding performance was significantly greater for the probe-matching vs. non-matching stimulus. (D) Cross-correlation between the task-relevant decoding time-series during the 100% and 75% conditions. The blue curve depicts the observed cross-correlation function while the black curve and grey shaded area depict a range of expected cross-correlation values simulated under the null hypothesis. (E) Peak decoding performance for the cue-matching disc during the 100% and 75% conditions. Error bars depict the 95% confidence interval of the mean.

Next, we examined location decoding performance during informative cue trials (Figure 4). During 100% reliable blocks, valid cues informed participants which disc would be probed with complete certainty. This condition is a direct replication of our earlier work (Ester et al., 2018) in which we found that valid postcues slowed or presented the gradual return to chance-level decoding performance seen during neutral trials. The current data replicate this finding (Figure 4A): during 100% valid trials decoding performance for the cue-matching disc remained at above-chance levels during the postcue display and into the probe display, while decoding performance for the cue-nonmatching disc quickly returned to chance levels following the appearance of the postcue. A qualitatively different pattern emerged during 75% reliable blocks (Figure 4B). Since invalid cues had no deleterious effect on participants’ recall errors (Figure 2B) or response times (Figure 2D), our analyses of decoding performance pooled across valid and invalid cue trials. Decoding performance for the cue-matching and cue-nonmatching discs returned to chance levels during postcue period, while decoding performance for the probe-matching disc (whether a valid or invalid trial) increased rapidly after the appearance of the probe display and remained at above-chance levels until after participants responded. Direct comparisons of cue- and probe-matching decoding performance during 100% and 75% blocks (Figure 4C) revealed that maximum decoding performance was reached significantly earlier during 100% blocks than 75% blocks (Figure 4D), even though averaged peak decoding performance was identical across these conditions (Figure 4E). Thus, the results of this analysis support the hypothesis that changes in behavioral priority affect the accessibility but not the strength of WM representations.

**Figure 4.**
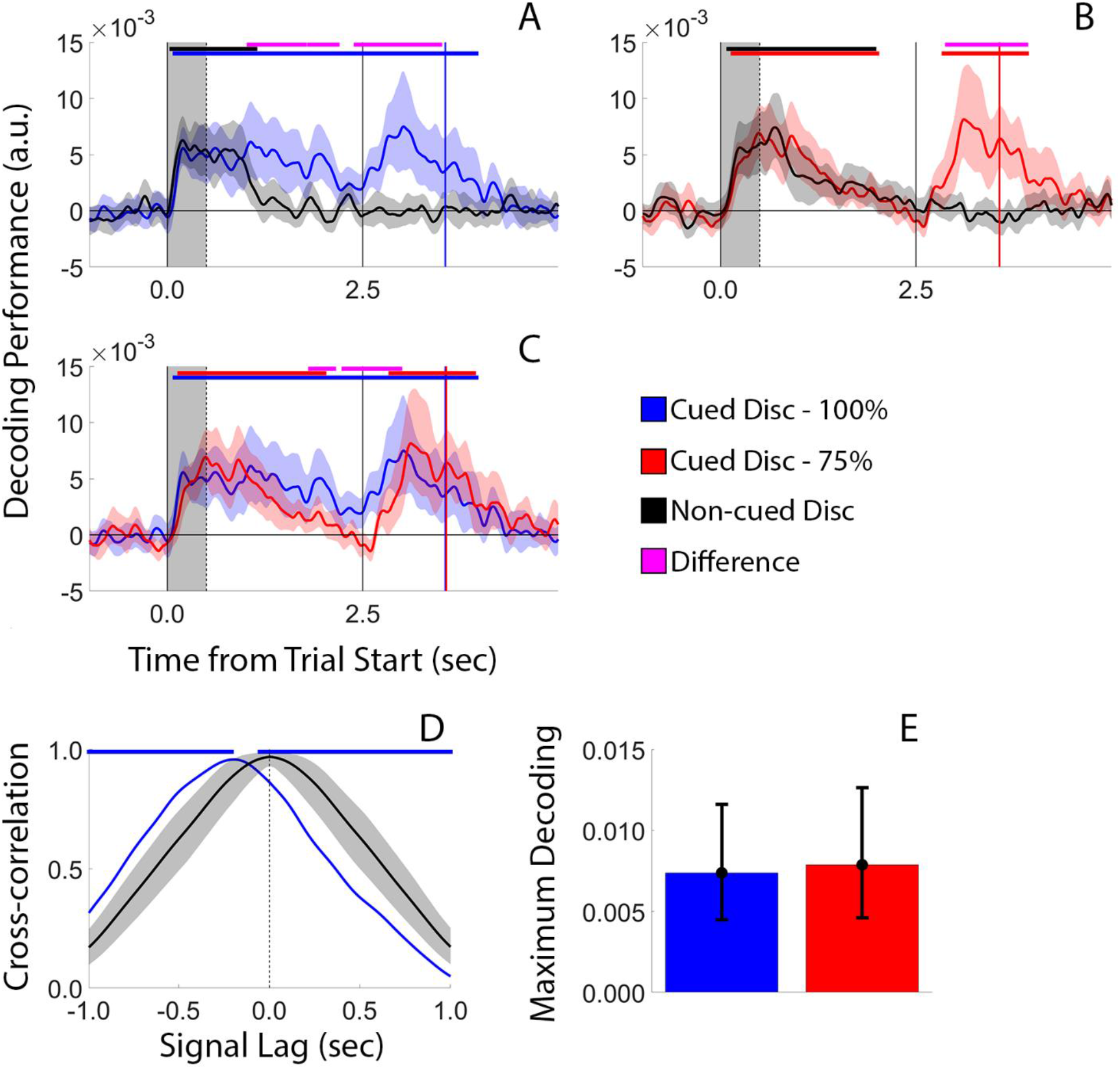
Decoding Performance During Informative Cue Trials. Conventions are identical to Figure 3.

### Experiment 2 – Retrocues

Experiment 2 was identical to Experiment 1, with the exception that informative postcues presented immediately after termination of the sample display were replaced by informative retrocues presented midway through the blank interval separating the sample and probe displays (see *Postcues vs. Retrocues*, Methods).

#### Behavioral Performance

Behavioral data from Experiment 2 were analyzed identically to Experiment 1. Participants’ recall errors during 100% blocks and 75% blocks are summarized in Figure 5A. Recall errors were significantly lower during valid compared to neutral trials during 100% blocks [t(32) = 2.637, p = 0.013, d = 0.179]. A repeated-measures analysis of variance (ANOVA) applied to recall errors during 75% blocks revealed a main effect of cue type (i.e., valid, invalid, neutral), [F(2,64) = 8.066, p = 0.0008, η^2^ = 0.201]. Cue effects – defined as the difference in recall errors between valid vs. neutral cues during 100% blocks and the difference in recall errors between valid and invalid vs. neutral cues during 75% blocks – are summarized in Figure 5C. Valid cues lowered recall errors by an equal amount during 100% and 75% valid blocks [M = 0.335° and 0.814°, respectively, t(32) = 1.918, p = 0.064], while invalid cues during 75% blocks incurred a significant performance cost compared to neutral trials [M = 1.086°, t(32) = 2.061, p = 0.048, d = 0.027].

**Figure 5.**
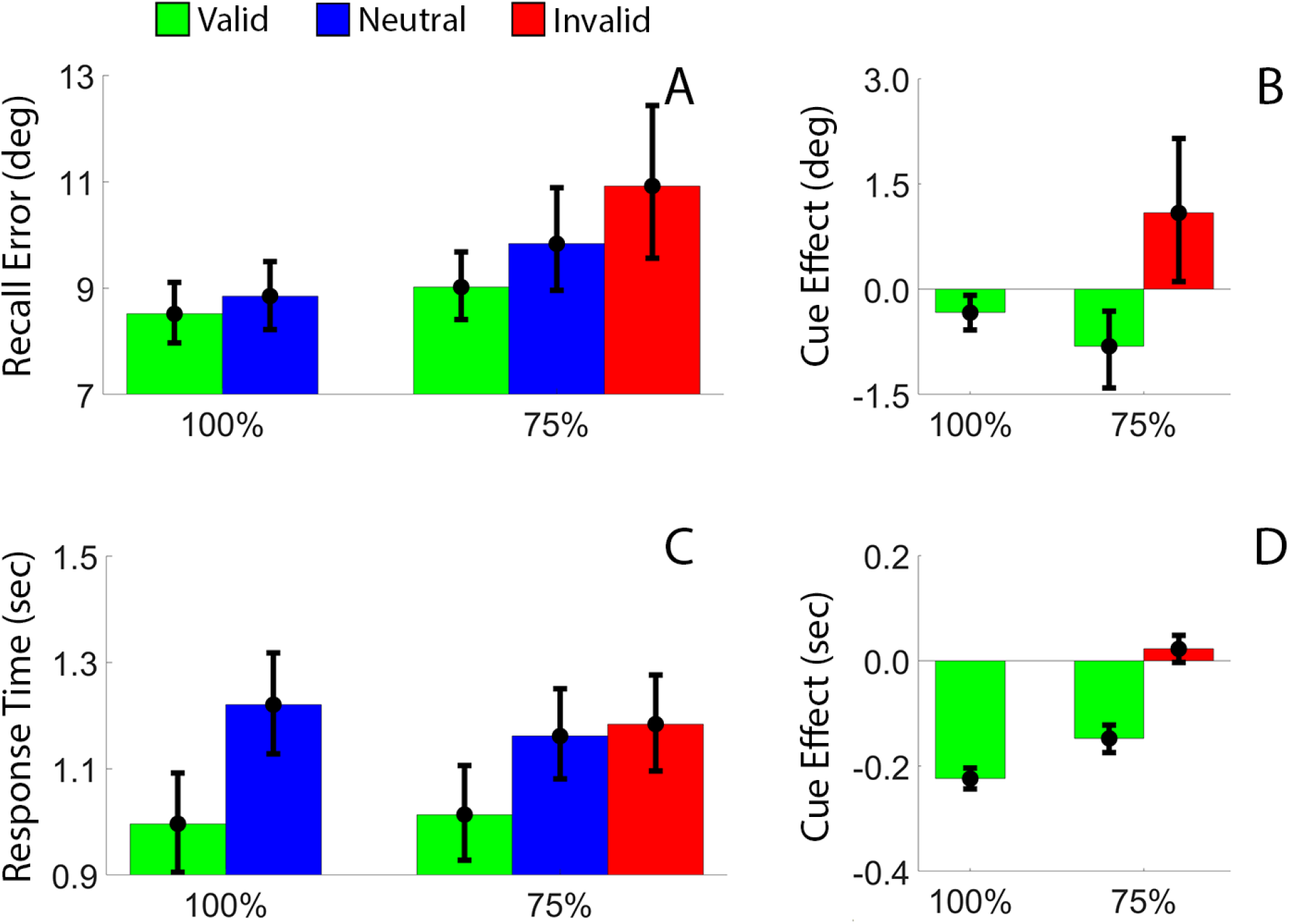
Behavioral Performance in Experiment 2. (A) Average absolute recall error as a function of cue type (valid, neutral, invalid) and cue reliability (100%, 75%). (B) Cue effects, defined as the difference between valid/invalid and neutral trials. (C) Average response times and (D) cue effects. Error bars depict the 95% confidence interval of the mean.

A complementary analysis of participants’ response times revealed a facilitatory effect of valid vs. neutral cues during 100% blocks [M = 0.997 vs. 1.220 sec, t(32) = 21.83, p = 8.66e-21, d = 0.804]. A repeated measures ANOVA applied to recall errors during 75% blocks revealed a significant effect of cue type (i.e., valid, invalid, neutral) [F(2,66) = 85.433, p < 1e-31; η^2^ = 0.728; Figure 5D]. Analyses of cue effects on response times revealed a greater benefit from valid cues during 100% vs. 75% blocks significantly larger effect of valid cues during 100% vs. 75% blocks [M = 224 vs. 148 ms, respectively; t(32) = 6.383, p = 3.597e-7, d = 1.093]. Invalid cues during 75% blocks did not incur a response time cost compared to neutral trials (M = 20.5 ms; t(32) = 1.702, p = 0.098, d = 0.084). Thus, valid retrocues improved participants’ recall errors and response times, and the magnitude of the response time benefit scaled with cue reliability (i.e., 100% vs. 75%).

#### EEG Decoding Performance

EEG data from Experiment 2 were analyzed in an identical way to those from Experiment 1. As in Experiment 1, we observed no effect of cue reliability on peak decoding accuracy or latency during neutral trials (Figure 6).

**Figure 6.**
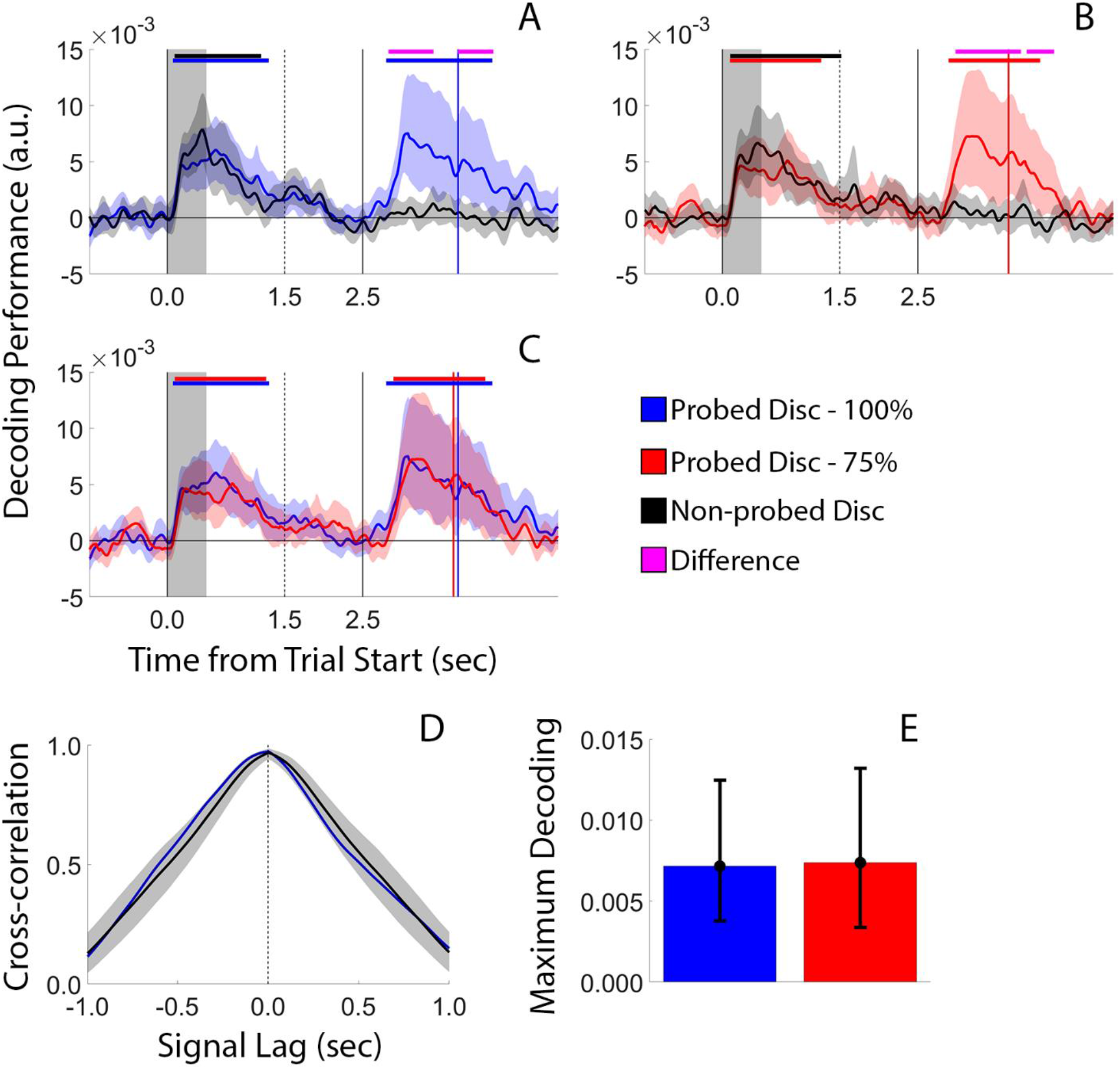
Decoding Performance During Neutral Trials of Experiment 2. Vertical lines at times 0.0, 1.5, and 2.5 sec mark the onset of sample, retrocue, and probe displays, respectively. All other conventions are identical to Figure 4.

Analyses of data from informative cue trials (Figure 7) were largely consistent with the findings of Experiment 1. During 100% blocks, decoding performance for the cue matching and non-matching discs increased rapidly following onset of the encoding display but returned to chance levels by the onset of the retrocue display 1.5 seconds later. Following the appearance of the retrocue, decoding performance for the cue-matching disc “recovered” to above-chance levels, replicating earlier findings showing cue-driven recovery in the quality of location-specific mnemonic representations (Sprague et al., 2016; Ester et al., 2018). Decoding performance for the cue-matching item remained at above-chance levels through the probe display and until shortly after participants’ responses. Conversely, decoding performance for the cue-nonmatching item remained at chance levels throughout the retrocue and probe displays.

**Figure 7.**
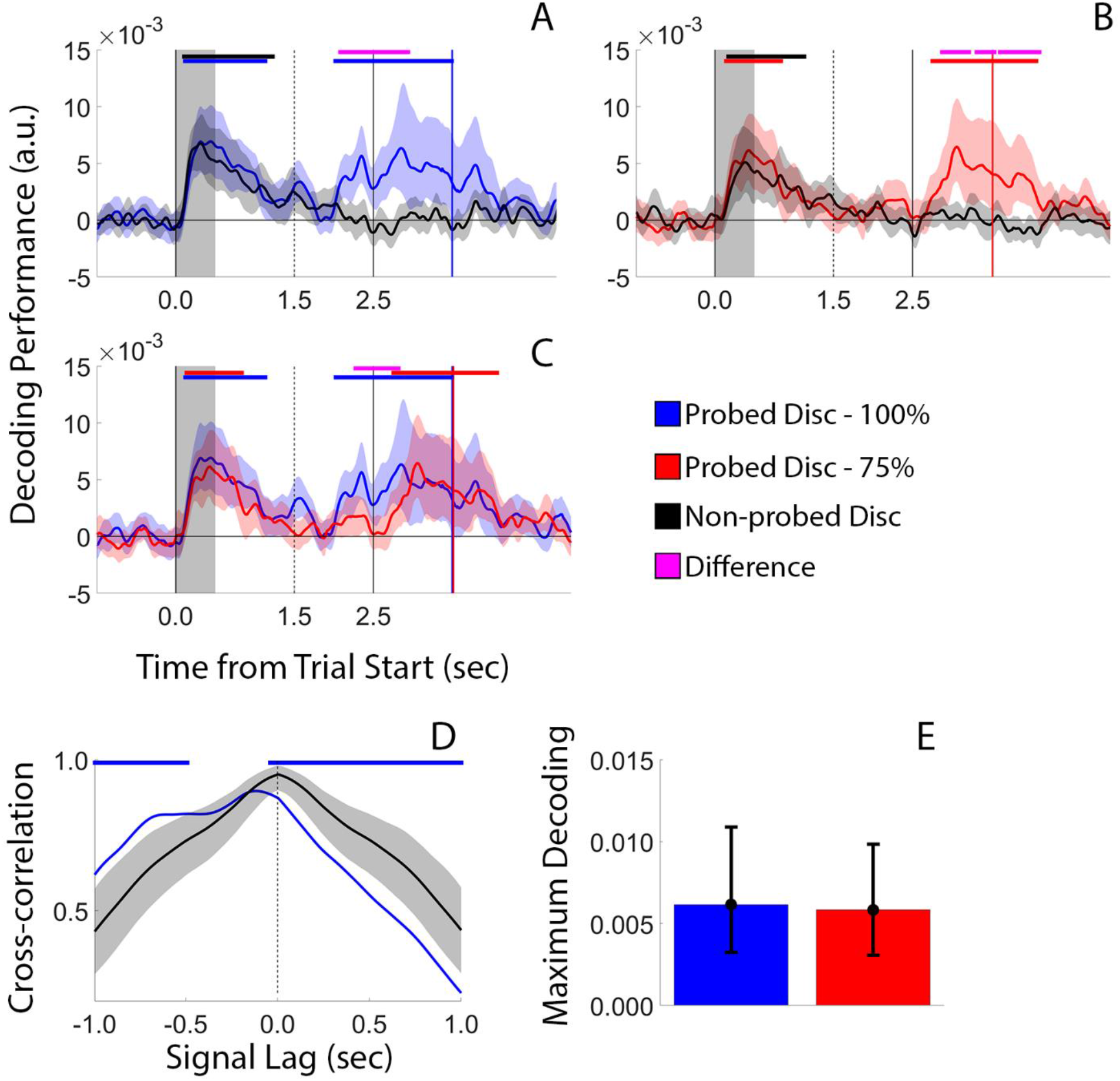
Decoding Performance During Informative Trials of Experiment 2. Conventions are identical to Figure 6.

An analysis of decoding performance during 75% blocks revealed a different pattern of findings (Figure 7B). Here, cue-matching and cue-nonmatching decoding performance remained indistinguishable from chance during the retrocue display, with cue/probe-matching decoding performance reaching above-chance levels only after the appearance of the probe display (as in Experiment 1, since invalid cues had no effect on participants’ recall errors or response times, we pooled data from valid and invalid trials to create the data shown in Figure 7B). Time courses of cue- and probe-matching decoding performance during 100% and 75% blocks are shown in Figure 7C. Comparisons of peak decoding latency (Figure 7D) revealed that maximum decoding performance was reached significantly earlier during 100% blocks relative to 75% blocks, although average peak decoding performance did not differ across these conditions. Thus, the findings of Experiment 2 are qualitatively identical to Experiment 1: changes in the priority of location-specific WM representations influenced the latency but not the magnitude of peak decoding performance.

#### Memory Retrieval or Response Preparation?

An alternative account of our findings holds that the differences in the onset timing of cue-locked above-chance decoding performance reflect response preparation rather than memory prioritization. We think this unlikely for several reasons. First, we note that the exact same physiological signal – total alpha power – was used for decoding throughout each trial, and that robust above-chance decoding performance was also observed during memory encoding (e.g., during the sample display and the early portion of the delay period) when participants had no way of knowing what item would be probed for report. Thus, proponents of a response preparation model must argue either that (a) above-chance decoding during memory encoding also acts as a form of response preparation (e.g., by encoding multiple different response affordances; Cisek & Kalaska, 2010), or that (b) above-chance decoding during memory encoding and retrieval – which, again, were computed using the exact same physiological signal – reflect WM and response preparation mechanisms, respectively. Either way, this argument would conflict with recent papers dissociating occipitoparietal alpha-band signals like those used for decoding in this study from response preparation and execution (e.g., van Ede et al., 2019; Boettcher et al., 2021; Ester & Weese, 2022). For example, van Ede et al. (2019) tracked occipitoparietal alpha power and frontocentral mu-alpha and mu-beta power while independently manipulating physical location (e.g., left vs. right visual field) of a to-be-recalled stimulus and the motor affordance (e.g., left vs. right hand) needed to perform recall. These authors found that occipitoparietal alpha power exclusively tracked the spatial position of the remembered item while frontocentral mu-alpha and -beta power exclusively tracked response demands. Thus, we think it unlikely that our findings can be explained by mechanisms of response preparation or execution.

Nevertheless, to obtain more traction on this issue, we examined the time-course of an EEG signal known to track response preparation and execution: lateralized frontocentral mu-alpha (∼8-13 Hz) and mu-beta (∼15-30 Hz) power. In this first phase of this analysis, we extracted total mu-alpha and -beta power from electrode site pairs C1/2 and C3/4. Our testing setup requiring all participants to respond with their right hand, so we computed mu-alpha and -beta lateralization by subtracting average power estimates from electrode sites C2 and C4 (i.e., ipsilateral to the response hand) from averaged power estimates from electrode sites C1 and C3 (i.e., contralateral to the response hand). We divided this difference by the sum of mu-alpha and -beta power over contralateral and ipsilateral sites to obtain a normalized (percentage) estimate of lateralization. During Experiment 1, mu-alpha and -beta lateralization steadily decreased (i.e., lower power over contralateral vs. ipsilateral electrode sites) over the interval separating the postcue and response displays, reaching a maximum shortly before the participant’s response (Figure 8). Importantly, neither the timing nor the peak magnitude of lateralization varied across cue conditions, i.e., 100% vs. 75%. A similar pattern was observed during Experiment 2 (Figure 9), with lateralization decreasing during the interval separating the retrocue and probe displays. To further test the response selection hypothesis, we also examined whether it was possible to decode the location of the cued/probed stimulus using frontocentral mu-alpha and mu-beta power. This analysis failed for both cue conditions (i.e., 100% vs. 75% valid) in Experiment 1 (Figure 10A-B) and Experiment 2 (Figure 10C-D). Thus, we argue that any observed differences in the timing or magnitude of location decoding performance are unlikely to reflect response preparation or execution. We describe the results of this analysis in the revised manuscript.

**Figure 8.**
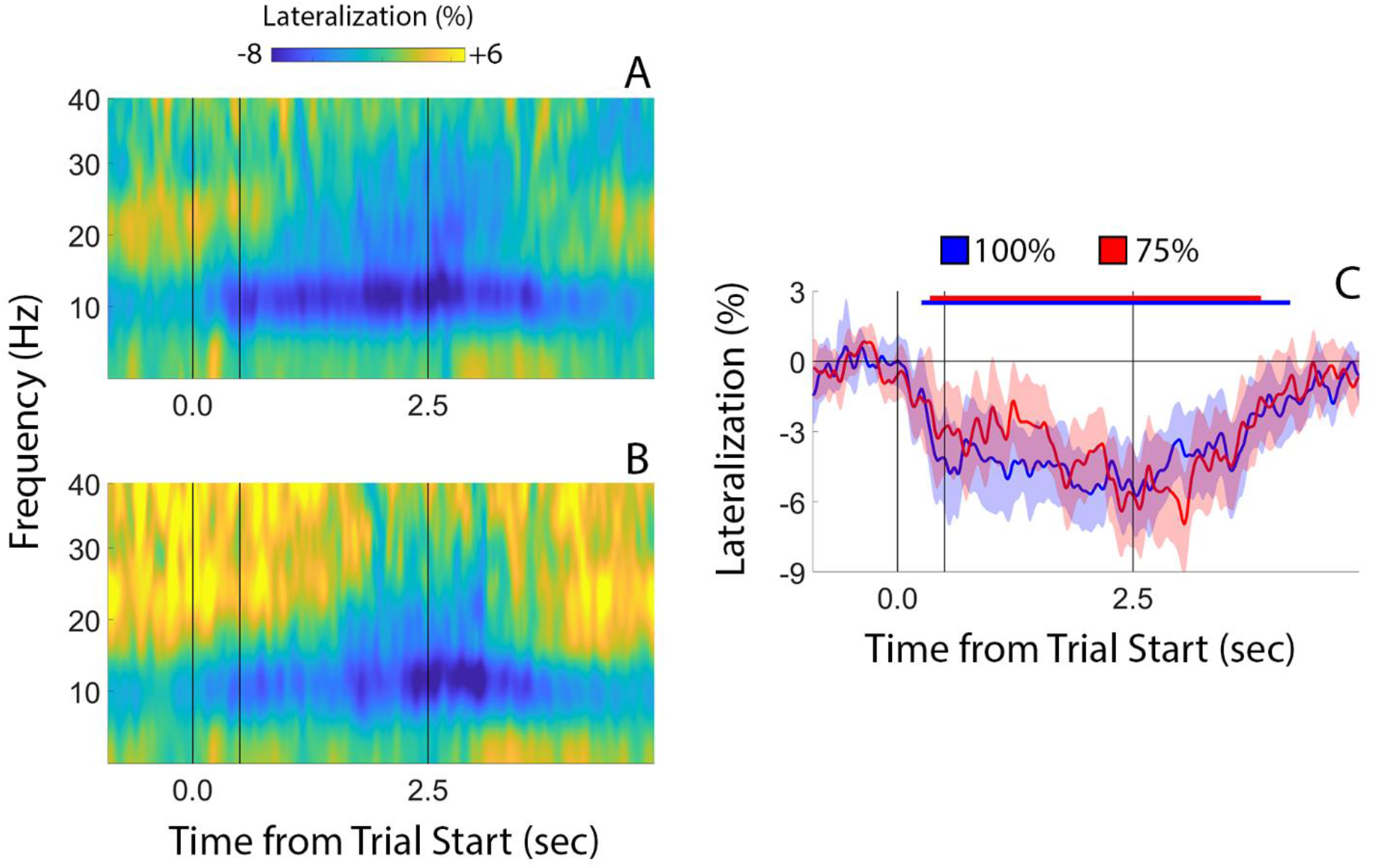
Frontocentral Signals Linked with Response Selection During Experiment 1. To test whether our core findings (e.g., Figs 4 and 7) could be explained by response selection, we tracked changes in lateralized frontocentral signals known to track response selection and execution. Analyses of lateralized frequency-specific activity revealed greater reductions in mu-alpha and beta-power over left – i.e., contralateral to the response hand – vs. right frontocentral electrode sites during the 100% task (A) and the 75% task (B). Next, we extracted, averaged, and plotted lateralized mu-alpha power (8-13 Hz) as a function of task (i.e., 100% vs. 75%; C). Although we observed robust reductions in mu-alpha power during both tasks, neither the timing or the peak magnitude of these effects differed across tasks. That we observed no differences in the timing or magnitude of an EEG signal known to track response preparation and execution suggests that the timing differences we observed in position decoding (e.g., Figs 4 and 7) cannot be explained by these factors.

**Figure 9.**
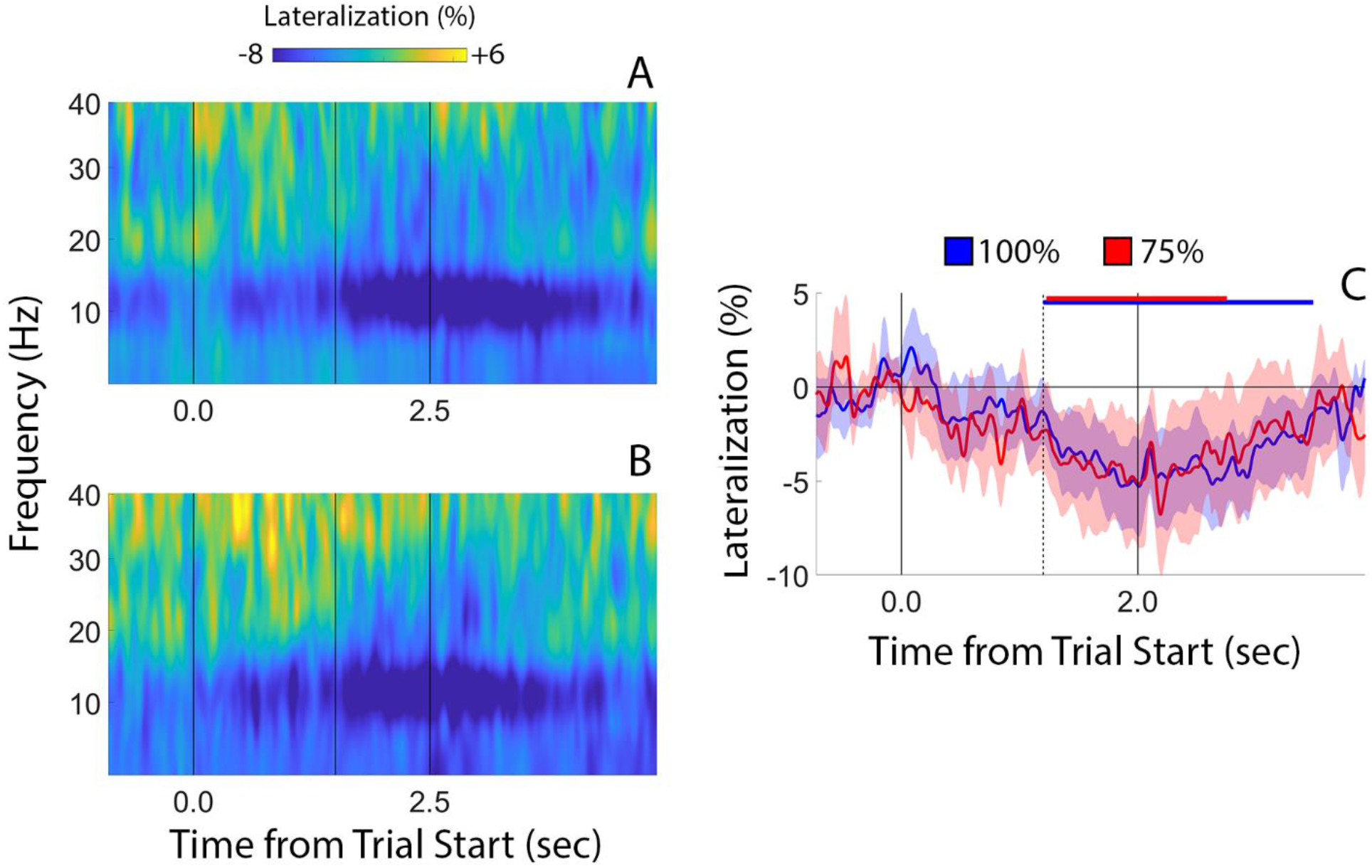
Frontocentral Signals Linked with Response Selection During Experiment 2. Conventions are identical to Figure 8.

**Figure 10.**
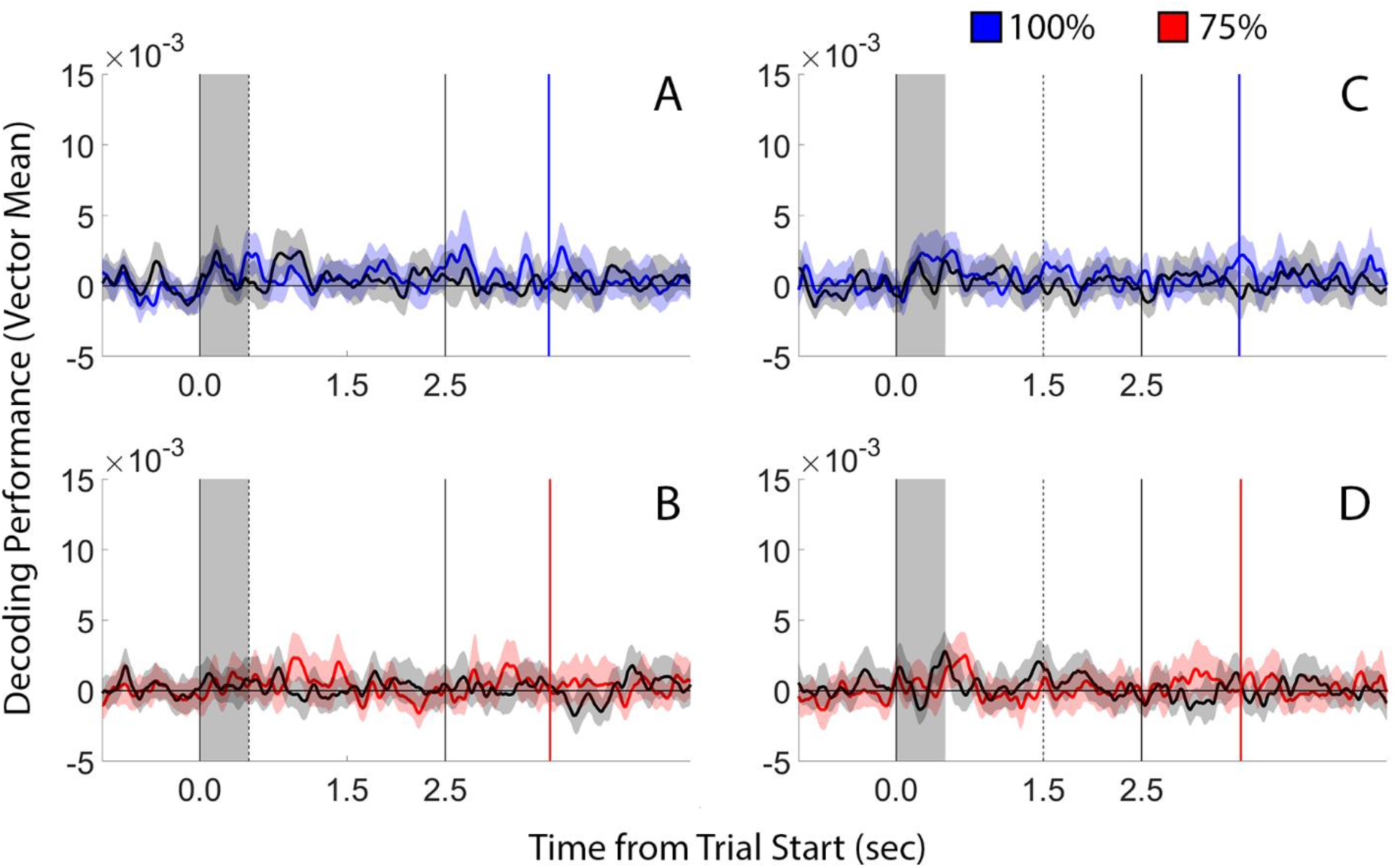
Position Decoding Performance Computed from Frontocentral Mu-alpha Power. As a further test of the response selection hypothesis, we attempted to decode the location of the probed (blue and red lines) and non-probed stimulus positions from frontocentral mu-alpha signals recording during informative cue trials in Experiment 1 (A-B) and Experiment 2 (C-D). This analysis did not support robust above-chance decoding of either the probed or non-probed position during either cue condition (i.e., 100% vs. 75%) or Experiment, providing further evidence against a response selection interpretation of our findings.

#### Alternative Explanations and Control Analyses

The data reported here suggest that access to cue-matching information is delayed when cue reliability is reduced (e.g., Figures 4D and 7D). One trivial possibility is that these findings are idiosyncratic to the parametric decoding approach we used or the alpha-band signals on which decoding performance was based. We tested these possibilities in complementary analyses.

First, we decoded the locations of the probed and non-probed discs from occipitoparietal alpha patterns using support vector machines (“one-versus-all” classification). Since stimuli could appear in eight possible locations, chance performance is 12.5%. The results of these analyses are summarized for Experiments 1 and 2 in Figures 11 and 12, respectively. Overall decoding performance computed using this approach was noisy (indeed, it was necessary to smooth the decoding time-series in Figures 11 and 12 with a 200 ms sliding window to identify clear trends in the data). Nevertheless, the overall pattern of findings obtained using this method was qualitatively similar to that obtained using our parametric decoding approach (e.g., compare Figures 4 and 11 and Figures 7 and 12). Critically, we again found no evidence for greater maximum decoding during the 100% vs. 75% cue reliability condition (Figures 11E and 12E).

**Figure 11.**
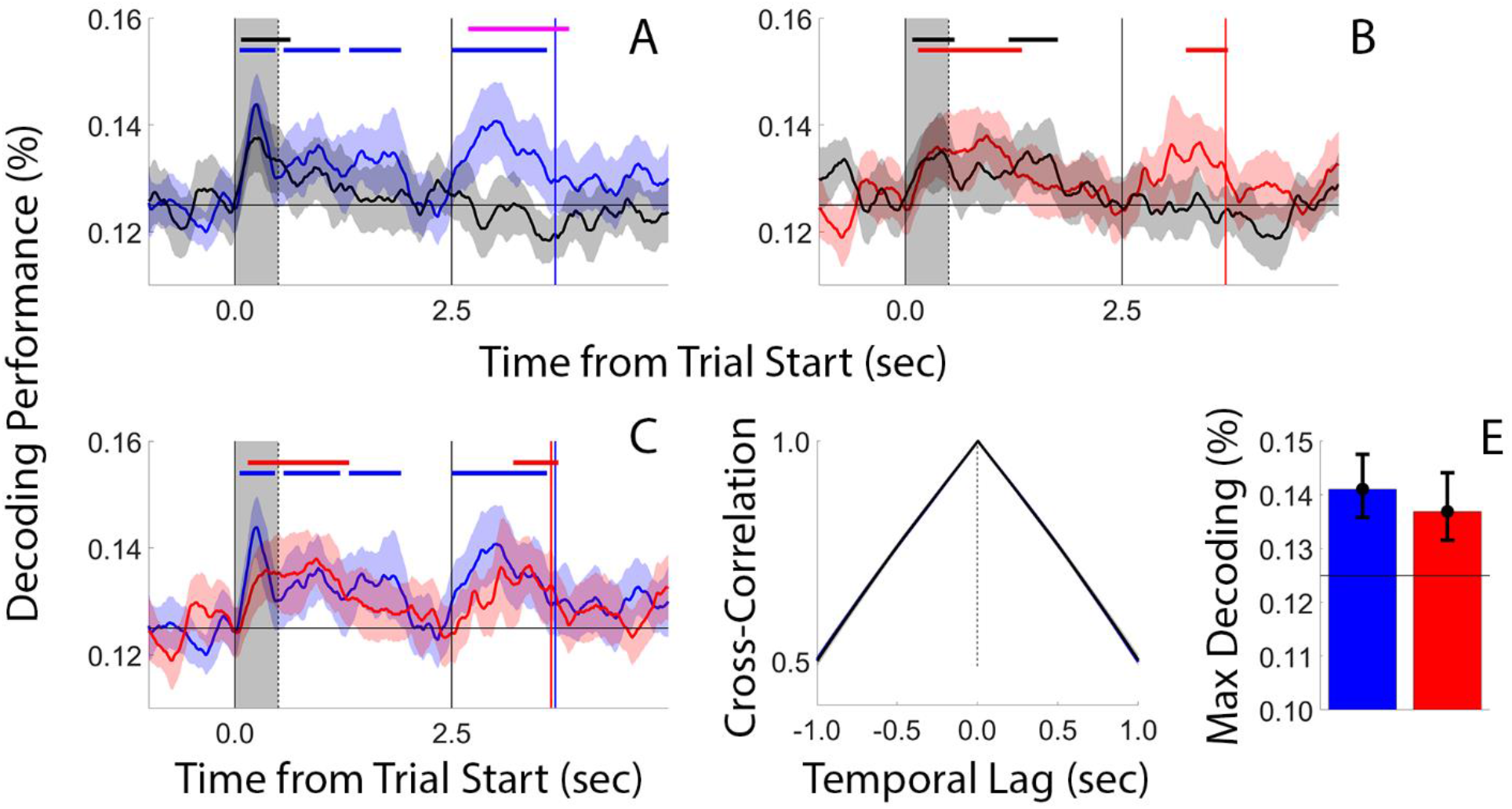
Experiment 1 Decoding Performance Computed Using Support Vector Classification. Conventions are identical to Figure 4.

**Figure 12.**
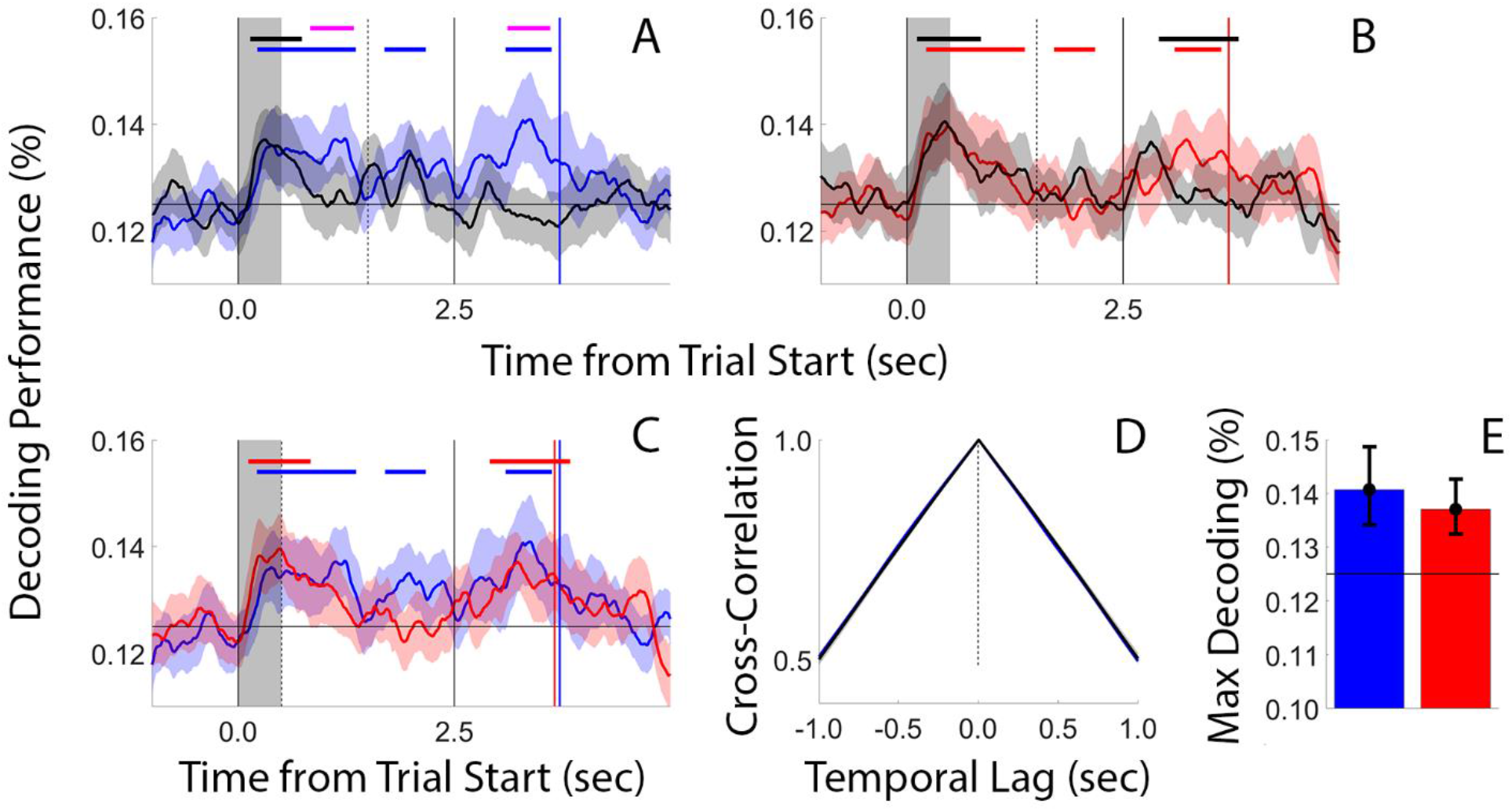
Experiment 2 Decoding Performance Computed Using Support Vector Classification. Conventions are identical to Figure 7.

Next, we asked whether position decoding performance was contingent on the use of spatiotemporal alpha power. On the one hand, some studies (e.g., Bae & Luck, 2018) have reported that alpha-band EEG signals uniquely index the positions of remembered stimuli while event-related potentials (ERPs) uniquely index the feature content of those memories. On the other hand, more recent studies (e.g., Barbosa et al., 2021) have shown that remembered orientations can be robustly decoded from patterns of occipitoparietal alpha-band activity. Since our experimental task required only memory for location, this may be a distinction without a difference. Nevertheless, we thought it prudent to establish that our core findings (Figs 4 and 7) generalize across different signal types. To this end, we used our parametric distance-based approach to decode the positions of the probed and non-probed positions during each experimental task and experiment using broadband EEG signals (i.e., voltages from 1-50 Hz). The results of these analyses were remarkably similar to the results obtained from decoding alpha-band signals for both Experiment 1 (Figure 13; compare with Figure 4) and Experiment 2 (Figure 14; compare with Figure 7). Thus, we are confident that our core findings (Figures 4 and 7) cannot be explained by idiosyncrasies unique to the alpha-band signal.

**Figure 13.**
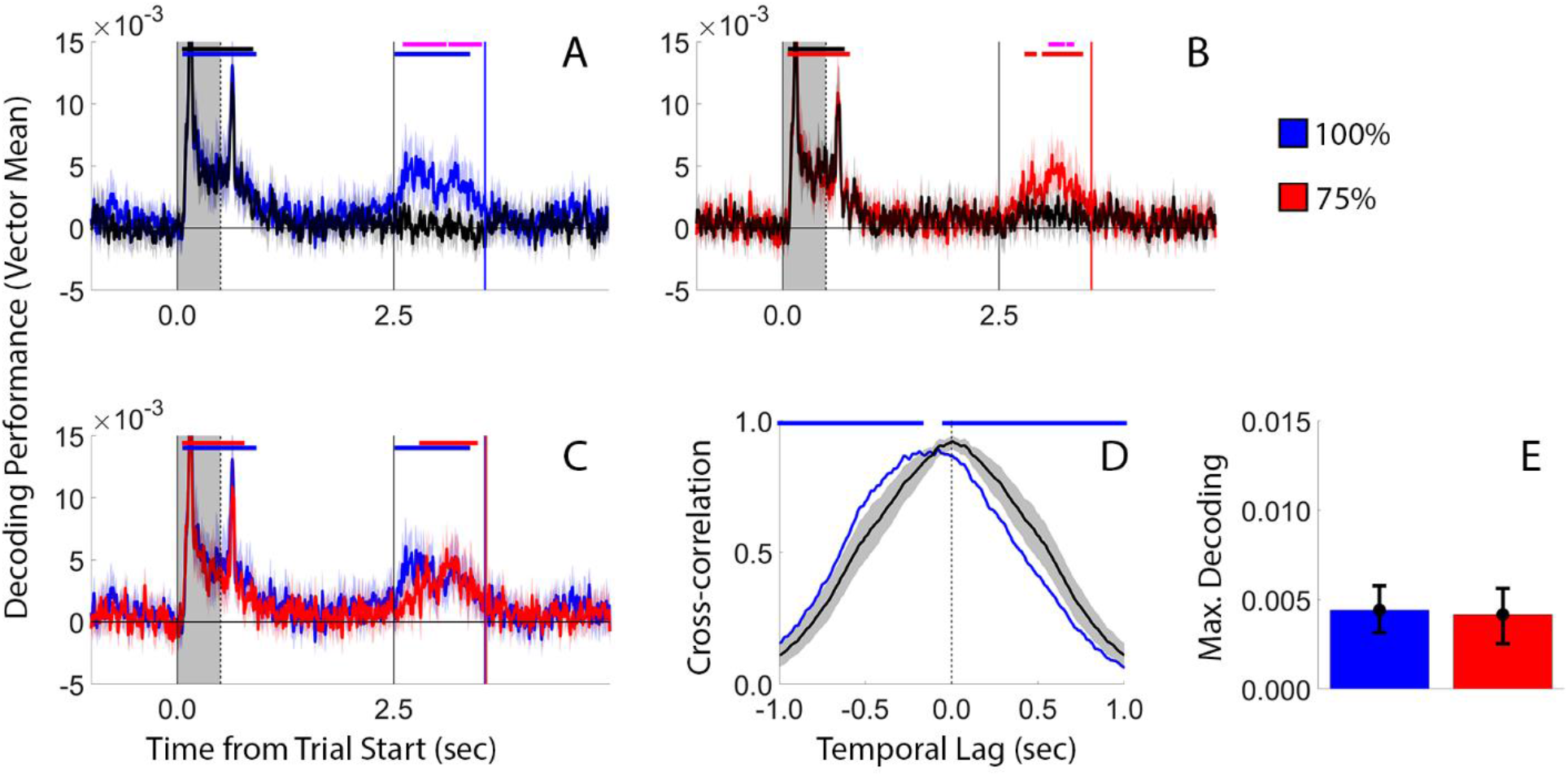
Experiment 1 Decoding Performance Computed from Broadband EEG Activity. Conventions are identical to Figure 4.

**Figure 14.**
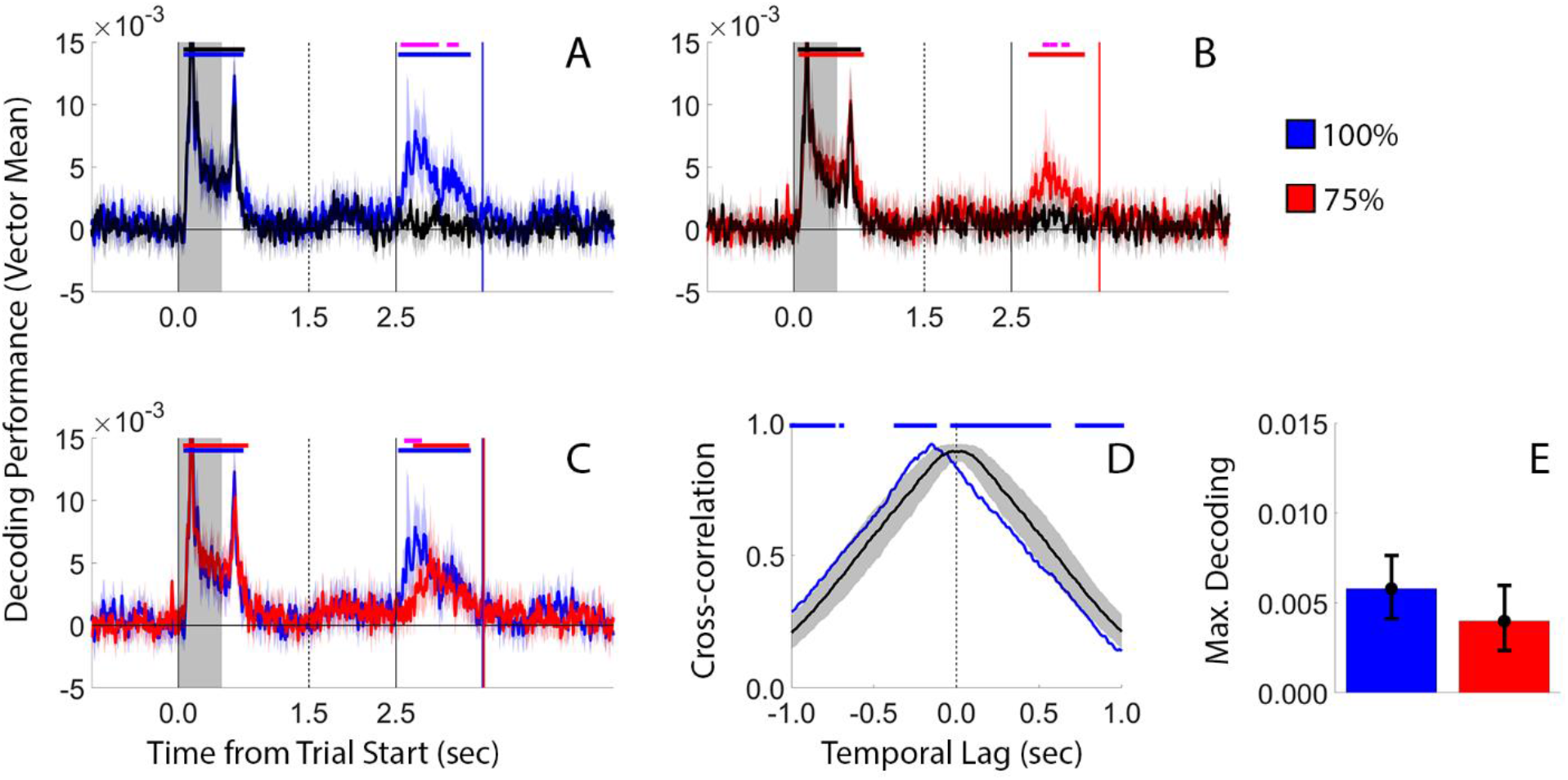
Experiment 2 Decoding Performance Computed from Broadband EEG Activity. Conventions are identical to Figure 7.

Next, we considered the possibility that our experimental manipulation of attentional priority was insufficient to detect changes in maximum decoding accuracy. Perhaps differences would be evident if we tested a larger range of cue reliabilities, e.g., comparing decoding performance across 100% vs. 60% reliability conditions or across 90% and 60% reliability conditions. Here, we note that our experiments also contained a 50% reliability condition: neutral trials. Thus, we performed direct comparisons between maximum decoding performance during informative cue and neutral cue trials from the 100% and 75% reliability tasks in each experiment. The results of these comparisons are summarized in Figure 15. Possible differences in maximum decoding performance were quantified via two-way repeated-measures analysis of variance (data from Experiment 1 and Experiment 2 were analyzed separately). When applied to data from Experiment 1, this analysis revealed neither a main effect of cue reliability (i.e., 100% vs. 75%; F(1,33) = 0.674, p = 0.417), a main effect of cue type (i.e., informative vs. neutral; F(1,33) = 2.933, p = 0.096), nor an interaction between these factors (F(1,33) = 1.159, p = 0.289). When applied to data from Experiment 2, this analysis also revealed neither a main effect of cue reliability [F(1,32) = 2.208, p = 0.147], a main effect of cue type [F(1,32) = 0.004, p = 0.948], nor an interaction between these factors [F(1,32) = 0.022, p = 0.882). Thus, no significant differences in maximum decoding accuracy were observed across a 50% reduction in cue reliability, supporting the view that the absence of task-level differences on this factor were not caused by a lack of sensitivity.

**Figure 15.**
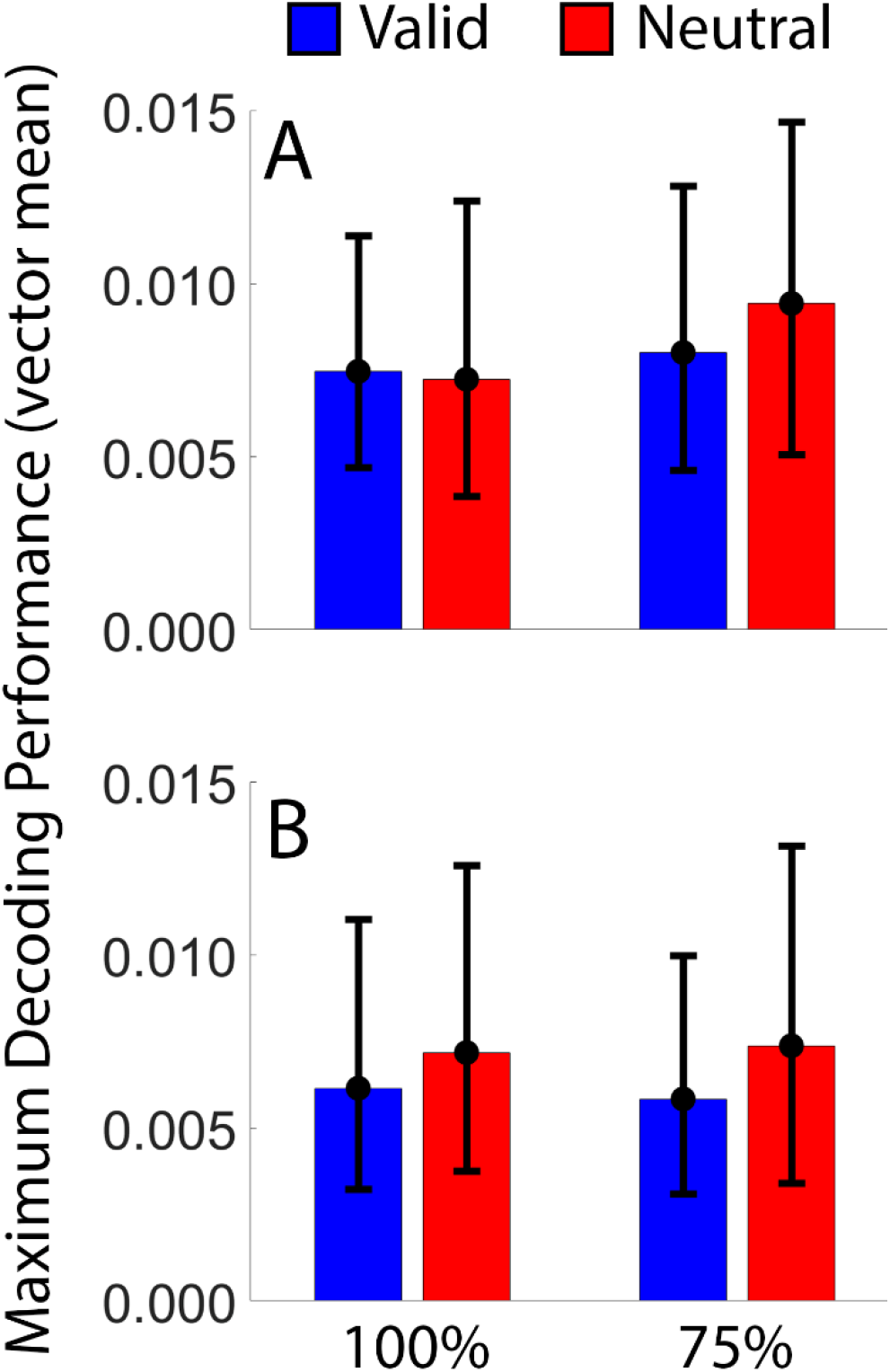
Comparison of Maximum Decoding Performance During Informative and Neutral Cue Trials During Experiment 1 (A) and Experiment 2 (B). Error bars depict the 95% confidence interval of the mean.

An alternative explanation for our findings holds that participants were simply more cautious or took more time to respond when cue reliability was fixed at 75% compared to when it was fixed at 100%, and that delays in the onset of above-chance decoding during the cue and probe displays reflect this caution rather than a delay in accessing the relevant WM representation. Analyses of participants response times do not support this possibility. Specifically, we reasoned that if participants were simply more cautious or slower in responding during 75% blocks, then their average response times during 75% valid trials should be significantly greater than during 100% valid trials. In fact, response times following valid cues were identical during 100% and 75% blocks in both Experiment 1 (M = 1058 vs. 1068 ms, respectively; t(33) = 0.579, p = 0.567; green bars in Figure 2C) and Experiment 2 (M = 996 vs. 1013 ms, respectively; t(32) = 0.768, p = 0.448; green bars in Figure 5C). We note, however, that cue effects (that is, the difference in response times across condition-specific neutral and valid trials, e.g., neural 100% vs. valid 100% trials) revealed significantly smaller response time benefits during the 100% vs. the 75% task. Thus, the findings reported here cannot be explained by general response caution or slowing during 75% vs. 100% blocks.

In both Experiments, cue-matching decoding performance during informative trials of 75% blocks fell to chance levels by the end of the memory period (Figure 4B and Figure 7B). This pattern is reminiscent of findings seen during neutral trials (Figures 3 and 6), raising the possibility that participants simply ignored the cues during 75% blocks. Once again, participants’ memory performance argues against this claim: recall errors and response times were significantly lower during valid vs. neutral trials during 75% blocks (Figure 2C and 5C). Nevertheless, to investigate the possibility that participants simply ignored informative cues during 75% valid blocks we undertook analyses comparing the time-courses of cue-matching decoding performance during 75% valid trials and 75% neutral trials. If participants indeed ignored informative cues, then the time-courses of decoding performance should be identical during informative and neutral trials. Conversely, if participants used informative cues to prioritize relevant information stored in memory, then maximum decoding performance should be reached earlier during informative vs. neutral trials. We tested these possibilities using the same cross-correlation analyses used to quantify differences in maximum decoding latency during 100% and 75% blocks, and the results are summarized in Figure 16. Maximum decoding latency was reached significantly earlier during informative vs. neutral trials of 75% blocks of Experiment 2 (Figures 16C and 16D), but not Experiment 1 (Figure 16A and 16B). These findings, coupled with analyses of participants’ memory performance, provide converging evidence against the possibility that delays in achieving maximum decoding performance during 75% relative to 100% blocks (Figure 4D and 7D) were caused by participants simply ignoring informative cues during 75% blocks.

**Figure 16.**
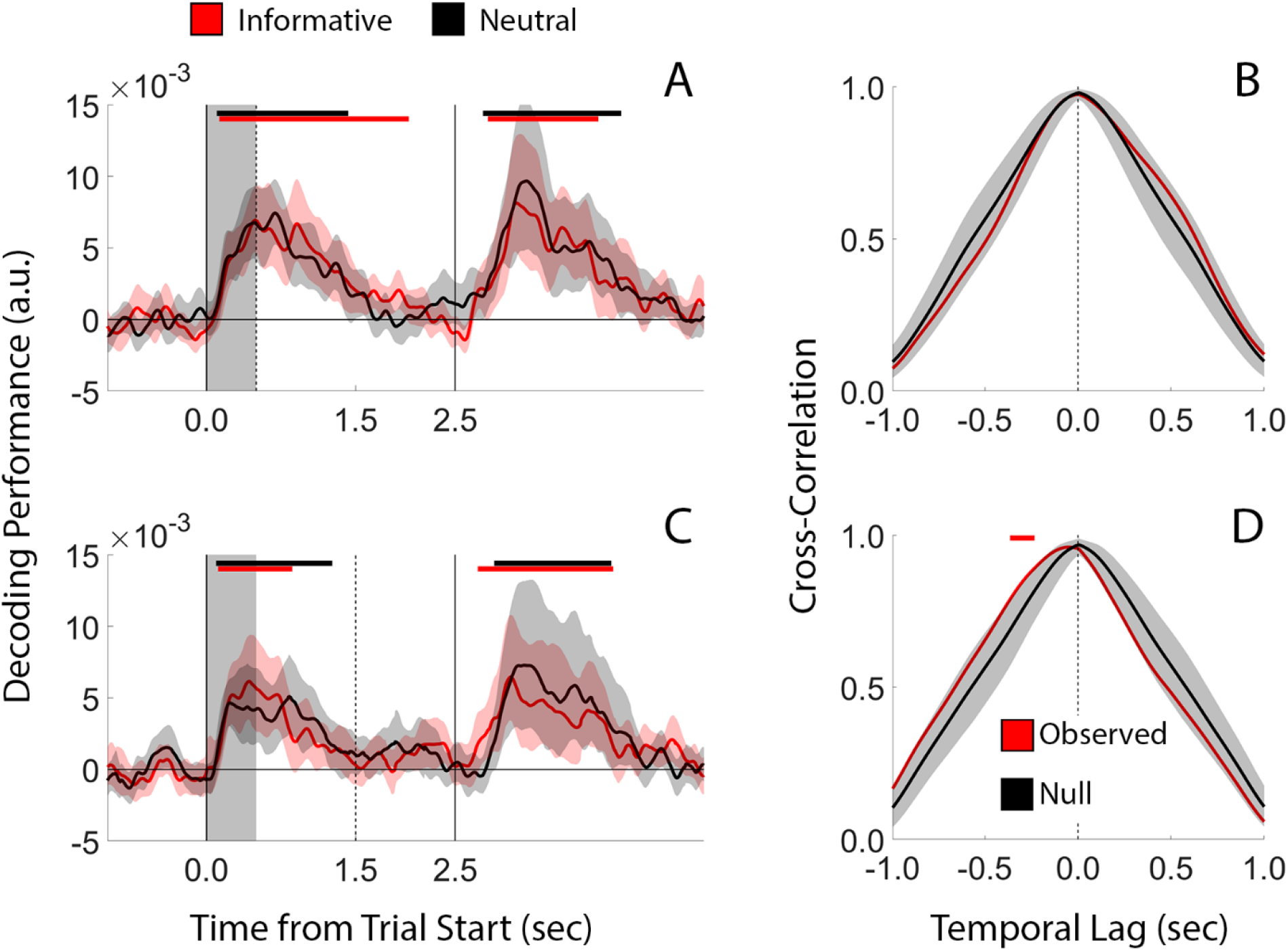
Comparisons of Above-Chance Decoding Latency on Informative and Neutral Trials during 75% blocks. (A) Overlay of above-chance decoding performance during informative and neutral cue trials of 75% blocks in Experiment 1. (B) Cross-correlation between probe-locked decoding performance during informative and neutral cue trials of 75% blocks in Experiment 1. Panels (C) and (D) are identical to panels (A) and (B) but use data from Experiment 2. Conventions are identical to those used in Figures 3-4 and 6-7.

## Discussion

Efficient behavioral selection requires rapid comparisons of sensory inputs with internal representations of motor affordances and goal states, and many of these comparisons take place in working memory (WM). Rapidly changing behavioral goals frequently require agents to assign (and re-assign) different levels of behavioral priority to items stored in WM. Prior studies utilizing retrospective cues suggest that humans can flexibly assign WM content with different levels of priority; for example, improvements in WM performance following an informative vs. uninformative retrospective cue scale positively with informative cue reliability (e.g., Berryhill et al., 2012; Shimi et al., 2014; Günseli et al., 2015; Günseli et al., 2019). Importantly, graded improvements in WM performance following a probabilistic retrocue could reflect (a) graded changes in the strength or quality of population-level, stimulus-specific neural patterns thought to mediate WM storage, (b) graded changes in how easily stimulus-specific neural patterns associated with high-(vs. low) priority items are accessed or “read-out” by downstream neural populations, or (c) some mixture or these options. Here, we leveraged the high temporal resolution of human EEG to adjudicate between these options. Specifically, we examined how graded changes in behavioral priority influenced our ability to decode a set of positions stored in WM. We reasoned that if priority-driven changes in memory performance are driven by the flexible allocation of attentional gain to items stored in memory, then population-level representations of high-priority items would be subject to larger gain modulations and thus easier to decode than low-priority items. Conversely, if priority-driven changes in memory performance are driven by changes in the accessibility of WM content, then it should be possible to decode the identity of high-priority items at an earlier time than low-priority items. Our data support the latter view. Specifically, we found no evidence suggesting that changes in behavioral priority – as manipulated via post- or retrocue reliability – influenced asymptotic decoding performance for cue-matching memoranda. Instead, the latency of asymptotic decoding performance was significantly earlier for high-priority vs. lower-priority items (Figures 4 and 7). Importantly, this effect could not be explained by mechanisms of response preparation or other nuisance factors (Figures 9-16). Thus, we conclude that changes in behavioral priority influence the accessibility but not the quality of stimulus-specific patterns of neural activity representing memoranda.

The term “priority” has been used to describe different phenomena in the WM literature (e.g., Riddle et al., 2020; Yu et al., 2020; Wan et al., 2022) and in visual neuroscience writ large (Rust & Cohen, 2022). Here, we use the term in the broadest sense to refer to cue-determined changes in the behavioral relevance of stimuli. This could occur, for example, when participants switch between sequentially reporting two items stored in WM (e.g., van Loon et al., 2018; Wan et al., 2022), when participants switch between preparing to report different items in WM following a retrospective cue (e.g., Lewis-Peacock et al., 2013; Rose et al., 2016), or when participants drop a subset of items stored in WM to focus on a different set of behaviorally relevant items stored in WM (Ester et al., 2018; Riddle et al., 2020; this study). In each of these examples, the sine qua non is an event signaling a change in the behavioral relevance of WM content. Importantly, what we term priority may reflect the operation of different mechanisms in different contexts. For example, in the case of sequentially reporting two items stored in WM, priority may refer to movement of memorized information from a latent to an active state. Likewise, in the case of switching between multiple potentially relevant WM items, priority may refer to the (internal) selection of likely task-relevant WM content. Research emphasizing transformations in the neural representations of WM content following changes in behavioral relevance (e.g., Panichello & Buschman, 2021; Bocincova et al., 2022; Li & Curtis, 2022) may help extirpate the use of colloquial terms like “selection”, “retrieval”, and “priority”.

Our analytic approach revealed evidence for an earlier active representation of cue-matching information during the probe epoch in the 100% vs. 75% condition (Figures 4 and 7). We speculate that this effect reflects differences in the time course of access to or retrieval of task-relevant WM content. Importantly the operation of retrieval processes need not be limited to the probe period where participants are required to make a behavioral response. Indeed, several lines of evidence suggest that human observers can flexibly access or retrieve information stored in WM following changes in behavioral relevance absent any response demands (e.g., Lewis-Peacock et al. 2013; Ester et al., 2018). Furthermore, concerns about comparing decoding time-courses during the 100% condition – where above-chance decoding performance is observed during the cue-to-probe interval – and the 75% condition – where above-chance decoding performance is observed only during the probe period – are somewhat mitigated by comparing decoding time-courses of above-chance decoding performance during 75% valid and neutral trials (Figure 16B). Here decoding performance is at chance levels during the cue-to-probe interval yet the onset of above-chance decoding performance occurs significantly earlier during 75% informative trials compared to neutral trials (even though, in both cases, the onset of the probe display coincides with the appearance of a perfectly informative response cue).

We have framed our discussion in terms of how changes in behavioral priority affect items that are likely to be probed for report, but some consideration should also be given to the effects of priority on items that are unlikely to be probed for report. For example, at least one previous study has reported that changes in behavioral priority affect the likelihood that cue-nonmatching items are attended and/or stored in WM (Günseli et al., 2019). The authors of this study used lateralized measures of covert spatial attention (alpha-band suppression; Klimesch, 2012) and WM storage (contralateral delay activity; Vogel & Machizawa, 2004) to show that cue-nonmatching items were less likely to be attended or retained in WM when cue reliability was low vs. high. Conversely, we found little evidence to suggest that changes in behavioral priority influenced neural patterns associated with memory for cue-nonmatching positions. However, a major difference between the current study and prior work is that we included 100% reliable cue blocks, which allowed participants to simply drop or forget cue-nonmatching information if they chose to do so. Thus, it is difficult to directly compare cue-nonmatching decoding performance across different levels of cue reliability (i.e., 100% vs. 75%). Additional traction on this issue could possibly be gained by comparing decoding performance for non-cued items during 75% blocks with decoding performance for both remembered items during neutral trials (akin to a 50% valid condition). However, we failed to observe above-chance decoding for the cue-nonmatching position during the delay period of 75% blocks or above-chance decoding for either remembered position during the delay period of neutral trials (e.g., Figures 3, 4B; 6, and 7B). Thus, additional research using a more fine-grained manipulation of cue reliability (e.g., 60%, 75%, 90%) will be needed to elucidate how changes in behavioral priority influence neural representations of cue-nonmatching items.

The current study utilized a mixture of postcues (Experiment 1) and retrocues (Experiment 2). Following earlier work (e.g., Souza & Oberauer, 2016) we use the term postcue to refer to any event informing which of a set of remembered item(s) will be tested that occurs immediately after encoding (including instances where an agent can apply this information to stimulus representations in sensory memory) and retrocue to refer to any informative event occurring after memory consolidation is complete. Prior evidence suggests that post- and retrocues engage separate but complementary mechanisms that promote storage of high-quality neural representations of memoranda. In an earlier study (Ester et al., 2018) we used an inverted encoding model to reconstruct location-specific representations of items stored in WM while presenting participants with neutral or perfectly informative postcues and retrocues. During neutral trials, the quality of reconstructed location-specific representations gradually decreased during WM storage. A perfectly valid postcue presented immediately after encoding eliminated this decrease, while a perfectly valid retrocue presented midway through storage partially reversed it. Data from 100% blocks in this study replicate these findings (e.g., Figure 4A; Figure 7A) while also establishing that assigning lower priority to items stored in memory reduces these effects (Figure 4B; Figure 7B).

In the absence of a perfectly informative postcue (i.e., Figure 4A), our ability to decode the location of the cue-matching or nonmatching disc fell to chance levels during the delay period (e.g., Figures 3A-C; Figure 4B; Figure 6; Figure 7B). This does not imply a loss of memory: participants still performed quite well during neutral trials despite no evidence for above-chance decoding during the delay period (Figures 3-4 and 6-7). One possibility is that the amount of location-specific information carried by induced alpha patterns that were used for decoding in this study gradually decreases over time. This, however, is difficult to reconcile with observations of robust above-chance location decoding during recall, as the probe display contained no additional information about the location of the to-be-reported disc (i.e., the color of the fixation point instructed participants which disc to recall, but gave no additional information about its position at the beginning of the trial). To account for this pattern, we speculate that during WM storage position-specific memory representations are gradually consolidated into a new format not indexed by alpha-band activity storage (for example, in an “activity-silent” synaptic network or in long-term memory; Rose et al., 2016; Sprague et al., 2016; Wolff et al., 2017; Masse, 2019; Barbosa et al., 2021; Beukers et al., 2021), and later retrieved from this format during memory recall.

WM can be conceptualized as a temporal bridge between fleeting sensory phenomena and possible actions. Recent theoretical conceptualizations of WM have begun to emphasize the action-oriented nature of this system (e.g., Olivers & Roelfsema, 2020; Heuer et al., 2020; van Ede & Nobre, 2022), and recent empirical findings suggest that behavioral (Ohl & Rolfs, 2020), circuit-level (Pho et al., 2018; Tang et al., 2020; Panichello & Buschman, 2021), and systems-level (Chatham et al., 2014; van Ede et al., 2019; Boettcher et al., 2021; Galero-Salas et al., 2021; Rac-Lubashevsky & Frank, 2021; Henderson et al., 2022) mechanisms of WM storage and action planning are tightly interwoven. In dynamic contexts where the future can take on several possibilities, the (potential) behavioral relevance of information stored in WM is often unknown. Thus, the likelihood that that any one piece of information stored in WM will become behaviorally relevant is best understood as a matter of probability rather than a certainty. From this perspective, a central purpose of WM may be to prepare for multiple potential futures, while mechanisms of internal attention act to select and prioritize relevant WM content as our predictions change or our uncertainties are reduced. The findings reported here are consistent with this view and further suggest that human observers can assign prospectively task-relevant representations different levels of priority that influences how quickly they can be retrieved and acted upon.

## Data Availability Statement

Stimulus presentation software, raw and preprocessed behavioral and EEG data files, and analytic software needed to generate each figure are freely available at https://osf.io/rg8h4/.

## Author Contributions

(using the Contributor Roles Taxonomy): E.F.E. – Conceptualization, Methodology, Software, Formal analysis, Investigation, Resources, Data curation, Writing – original draft, Supervision, Funding acquisition. P.P. – Methodology, Investigation, Writing – review & editing.

